# Continuous transcriptome analysis reveals novel patterns of early gene expression in *Drosophila* embryos

**DOI:** 10.1101/2022.09.26.509035

**Authors:** J. Eduardo Pérez-Mojica, Lennart Enders, Joseph Walsh, Kin H. Lau, Adelheid Lempradl

## Abstract

Early organismal development consists of transformative events that lay the foundation for body formation and long-term phenotype. Despite this understanding, the rapid progression of events and the limited material available are major barriers to studying the earliest stages. The size and accessibility of *Drosophila* embryos overcome some of these limitations, and several studies characterizing early transcriptional events have been reported. Unfortunately, manual embryo staging, and elaborate protocols make the techniques employed in these studies prone to human and technical errors and incompatible with routine laboratory use. Herein, we present a straight-forward and operationally simple methodology for studying the early transcription (≤3 hours) in *Drosophila*. This method relies on single-embryo RNA-sequencing and transcriptome ordering along a developmental trajectory (pseudo-time), thereby avoiding the need for the staging of the embryos. The obtained high-resolution and time-sensitive mRNA expression profiles uncovered the exact onset of transcription and degradation of transcripts and revealed an earlier transcription start for several zygotic genes. In addition, degradation patterns suggest that maternal mRNA decay is independent of mRNA levels. By classifying each embryo as male or female, we were also able to study sex-biased transcription between the beginning of zygotic transcription to gastrulation and identified 120 differentially expressed mRNAs. Using sex-specific transcription signatures, embryos can be sexed directly, eliminating the need for Y-chromosome genotyping. Herein, we report an accessible, single-embryo sequencing approach for high-resolution, time-sensitive transcriptome analysis. Our data provide an unparalleled resolution of gene expression during early development and enhance the current understanding of early transcriptional processes.

## INTRODUCTION

In many animal species, the zygote relies on maternally deposited transcripts to progress through the earliest stages of development (Vastenhouw et al. 2019). It is not until later that the zygote takes control of its own development; a process referred to as the maternal-to-zygotic transition. An important element of this transition is the start of transcription from the zygotic genome, also referred to as zygotic genome activation (ZGA). Previous studies have shown that the ZGA is a progressive event. It starts with the transcription of just a handful of genes (minor ZGA) and increases to hundreds of genes shortly thereafter (major ZGA) (Lott et al. 2011; Sandler and Stathopoulos 2016; Kwasnieski et al. 2019).

In *Drosophila melanogaster*, it is generally accepted that *sisterless A* (*sis A*) and *snail* (*sna*) are transcribed early in development during nuclear cycle (NC) 8 (Erickson and Cline 1993; Pritchard and Schubiger 1996). Evidence suggests that *scute* (*sc*) is an additional early transcribed gene, but reports differ on the timing of transcriptional onset (Erickson and Cline 1993; Parkhurst et al. 1993; Deshpande et al. 1995; Ali-Murthy et al. 2013). The total number of zygotic transcribed genes reported to be expressed by NC 9 is 20 and increases to 63 by the end of NC 10 (Kwasnieski et al. 2019). This so-called minor wave of transcription coincides with other important developmental processes, such as the migration of nuclei to the posterior pole of the embryo and the generation of pole cells (Foe and Alberts 1983). With the onset of the minor ZGA, nuclear cycles become progressively longer. While the first 8 NCs on average take eight minutes, the duration continuously increases until it reaches 65 min at NC 14 (Foe and Alberts 1983). NC 14 marks the beginning of the major ZGA and the number of genes transcribed increases significantly to 3,540 (Kwasnieski et al. 2019). The major ZGA is accompanied by important developmental processes such as cellularization, first gastrulation movements, end of synchronous nuclear divisions, and the start of dosage compensation by Male-Specific Lethal (MSL) complex (Lott et al. 2011; Farrell and O’Farrell 2014). Of note, sex-specific transcription is observed as early as NC 11 (Lott et al. 2011). The exact onset and sequence of transcriptional events leading up to this differential gene expression remains poorly understood. In parallel to ZGA maternal transcripts are being degraded in an organized manner. This process of clearing of maternally deposited mRNAs is essential for proper development. It is important to note that the degradation of certain maternal transcripts occurs even in unfertilized eggs (Thomsen et al. 2010).

The limited size of embryos and rapid progression of developmental processes make a quantitative assessment of transcriptional events challenging. Previous studies have investigated the timing, extent, and sex-specificity of the ZGA using different methods such as RNA radioactive (Edgar and Schubiger 1986) or metabolic labeling (Kwasnieski et al. 2019), in situ hybridization (Tomancak et al. 2002; Hammonds et al. 2013), RNA sequencing (RNA-seq) (Lott et al. 2011, 2014), qPCR-based experiments (Ali-Murthy et al. 2013), and direct count of mRNA molecules (Sung et al. 2013; Sandler and Stathopoulos 2016). Notably, all the different methods rely on the meticulous staging of embryos, which is time consuming, error prone, and requires a well-optimized protocol to guarantee the fast collection of embryos when working with fresh samples. Currently, the only available option to avoid manual embryo staging is to rely on short egg laying times and incubation to the desired developmental stages. The results of these studies, however, can be biased due to the rate of egg fertilization, regularity of oviposition, and embryo withholding. The latter has been shown to differ for more than 10 hours in some *Drosophila* species (Markow et al. 2009). The technical difficulties of existing protocols have led to inconsistencies between findings from different laboratories. For instance, more than half of the transcripts assigned to the minor ZGA in one study (Ali-Murthy et al. 2013) were likely due to the contamination of a sample with an older embryo (Kwasnieski et al. 2019).

To address these technical limitations and ensure increased data reproducibility, we developed a single-embryo RNA-seq method to measure zygotic transcripts on a continuous time scale. Using an analysis pipeline designed for single-cell RNA-seq, we utilize the transcriptome to determine the biological age and sex for each embryo, eliminating human and technical errors introduced by visual staging. The data produced using this method can be corroborated through comparison with published data and provide the first continuous timeline of transcript levels during early development (≤ 3 hours) in *Drosophila melanogaster*.

## RESULTS

### Single-embryo RNA-sequencing

In order to study early embryonic transcription in a continuous manner, we performed single embryo RNA-seq on a total of 192 embryos. The embryos were collected from two different cages in three consecutive one hour (h) time intervals and allowed to develop further for 0, 1 or 2 hours. This resulted in an approximate collection time window of 10 minutes to 3h. RNA was isolated from individual embryos to perform single-embryo RNA-seq using a modified CelSeq2 protocol (Hashimshony et al. 2016; Sagar et al. 2018). The sequencing data were analyzed using the RaceID3/StemID2 single-cell analysis tool (Fig. 1A) (Herman et al. 2018). Embryos with less than 250,000 reads were excluded from the analysis, leaving 122 embryos for final analysis. In total, we identified 9777 genes with ≥ 3 unique molecular identifier (UMI) corrected read counts in ≥ 5 embryos. Unsupervised *k*-medoids clustering of our data, according to transcriptome similarities, resulted in 14 clusters (Fig. 1B). Dimensionality reduction of the single-embryo RNA-seq data using t-distributed stochastic neighbor embedding (t-SNE) produced a map where all samples assembled in a linear pattern (Fig. 1B). A similar arrangement was confirmed by other dimensionality reduction methods like a Fruchterman-Reingold force directed layout or principal component analysis (PCA) (Supplemental_Fig_S1A,B.pdf). Because mated females can lay unfertilized eggs, which would compromise our analysis, we used the expression of previously reported early transcribed genes (*screw* (*scw*), *scute* (*sc*), and *escargot* (*esg*)) to identify and exclude such embryos (Supplemental_Fig_S1C,D,E.pdf). The number of unfertilized eggs (n = 5) in our dataset matches the number expected, given the 0.95 fertility rate that was measured on the same day as sample collection (Supplemental_Fig_S1F.pdf). Analyzing fertilized embryos only resulted in a layout similar to the one observed for all samples (117 embryos, Fig. 1C). We then used StemID2, an algorithm developed for the derivation of differentiation trajectories in single-cell data, to generate a lineage tree object, where each embryo receives a relative coordinate on the inferred inter-cluster links according to their transcriptome. Fig. 1D shows the projection of the embryos onto a minimum spanning tree of a predicted differentiation trajectory. This ordering of embryos along a computed developmental trajectory is also called pseudo-time. Both t-SNE and pseudo-time analysis show that there is no cage batch effect in either analysis (Supplemental_Fig_S1G,H.pdf). For our data, pseudo-time encompasses the time it takes from 10 minutes after fertilization (delay necessitated by sample processing time) to the developmental stage represented by the last embryo on the spanning tree. We subsequently compared this computationally derived pseudo-time with our 3 sample collection time intervals. Even though embryos were collected in three defined one-hour intervals, their position in pseudo-time was not restricted to their respective collection time window (Fig. 1E, Table 1). These results confirm the propensity of mated females to withhold fertilized eggs for extended periods of time. To avoid contamination of samples by withheld embryos, published related methods currently rely on the laborious and error prone processes of hand staging the developing embryos. Our method enables us to identify these withheld embryos and assign them to their correct developmental time without the elaborate process of visually staging the embryos. Together, these results show that we successfully developed a single embryo RNA-seq protocol and analysis pipeline without the use of elaborate labeling protocols or staging techniques.

**Figure 1.**
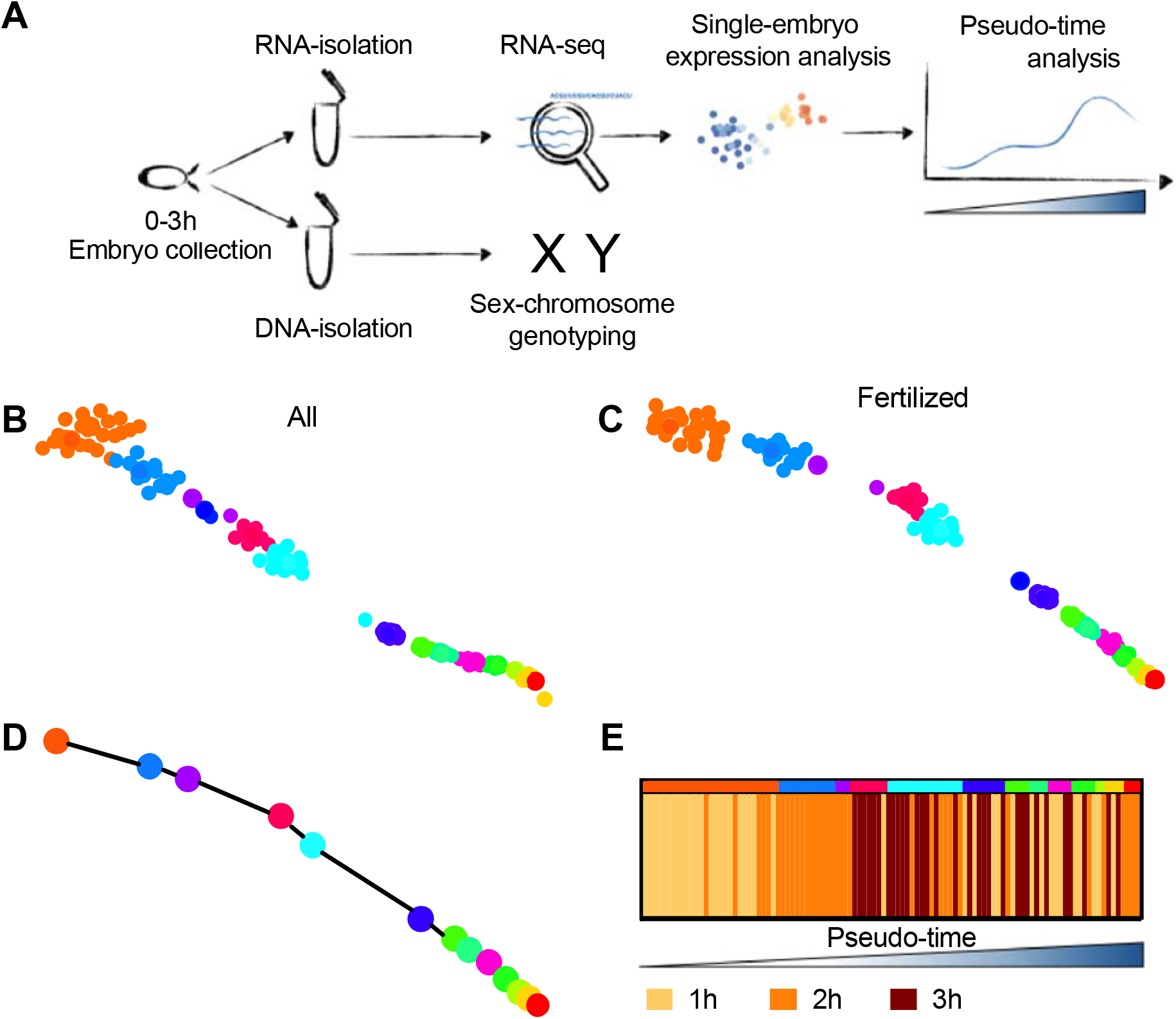
Single-embryo RNA-sequencing approach to bioinformatically identify developmental age. (**A**) Schematics of methodology: single embryos are collected in 1-hour (h) intervals and aged up to 3h. RNA and DNA are isolated from the same single embryos. DNA is used for genotyping the X and Y-chromosome, while RNA is processed using a modified CEL-Seq2 protocol to determine embryo age. (**B**) t-SNE before (n=122) and **(C)** after the removal of unfertilized eggs (n=117) with clusters identified by k-medoids clustering indicated by different colors. (**D**) Lineage analysis by StemID2/FateID3 identified a single trajectory for all clusters (with n > 1 embryos) resulting in the ordering of embryos along a pseudo-time axis according to their age. (**E**) Comparison of the pseudo-time order with the actual collection time intervals. Ascending pseudo-time (embryo age) from left to right, colors in top bar indicate clusters from (C).

**Table 1.** Number of embryos in each quartile of the pseudo-time by collection time

|  |  | Number of embryos in<br>collection time-intervals |  |  |
| --- | --- | --- | --- | --- |
|  |  | 0-1 h | 1-2 h | 2-3 h |
| Pseudo-time<br>quartiles (Q) | Q1 | 22 | 4 | 0 |
|  | Q2 | 2 | 17 | 7 |
|  | Q3 | 5 | 7 | 14 |
|  | Q4 | 10 | 6 | 10 |

### Single-embryo sequencing and pseudo-time analysis allow for the continuous assessment of transcription profiles during early embryogenesis

Next, to investigate the expression of genes across early development, we focused our analysis on approximately 3h old embryos (84 embryos). This analysis revealed 9 stable clusters of embryos based on their transcription (Fig. 2A,B) and provided a more refined look at the developmental trajectory compared to our previous analysis that included all embryos (Fig. 1B). We next sought to determine if the computationally derived pseudo-time was in agreement with the biological age of the embryos. To this end, we assessed each embryo for the expression of genes that have been previously reported to be activated during the minor (Kwasnieski et al. 2019) or major (Sandler and Stathopoulos 2016) wave of ZGA (Supplemental_Table_S1.pdf). Plotting the combined expression of these genes onto our 2-dimensional layout (Fig. 2C,D) and along the pseudo-time scale (Fig. 2E,F) reveals that these two major transcriptional events coincide with the increased distance in our t-SNE map between clusters 1 and 2 and 4 and 5. This is expected as gaps like this indicate major transcriptional shifts and therefore validates our computational approach.

**Figure 2.**
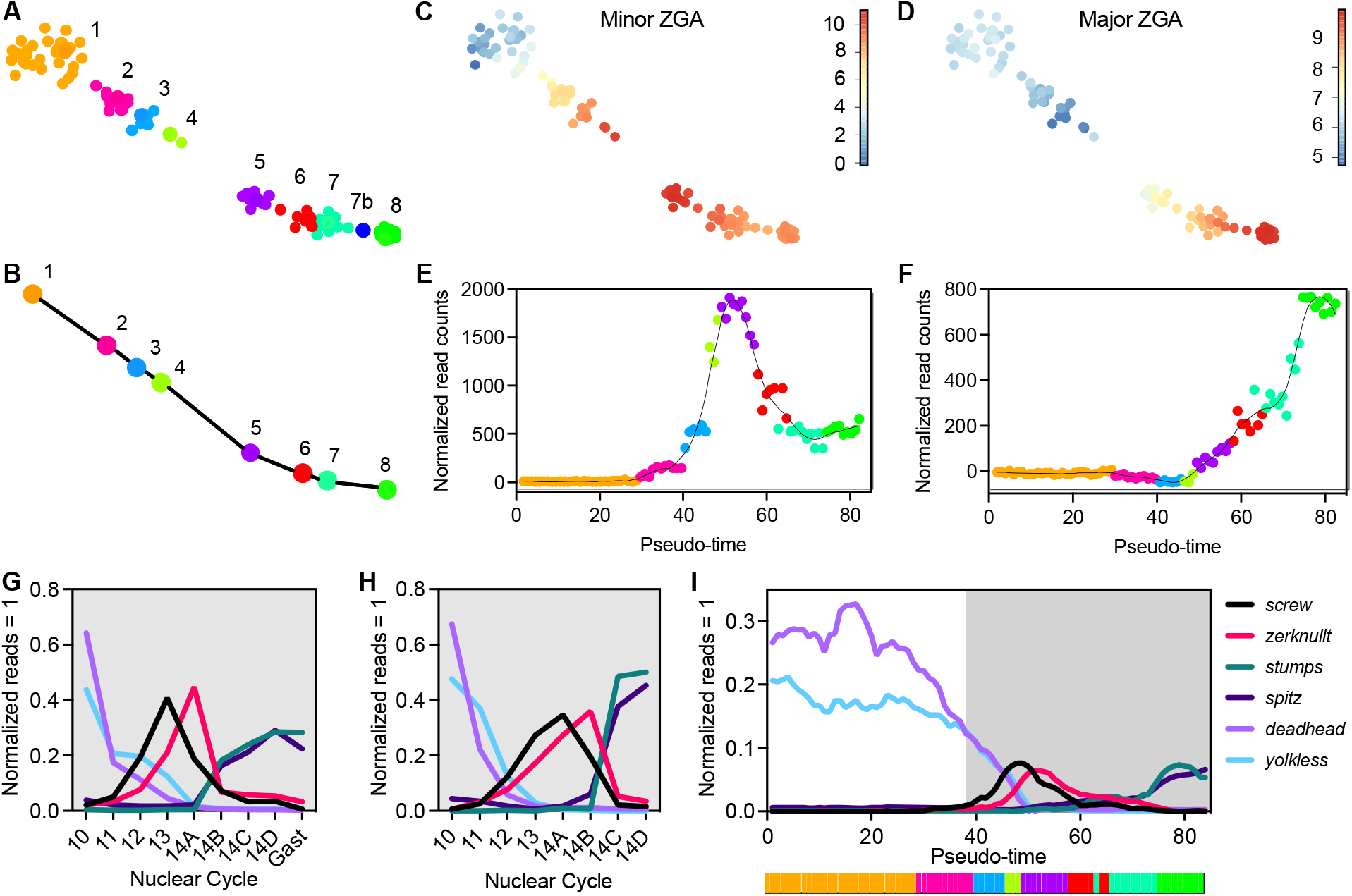
The continuous sequence of the zygotic genome activation (ZGA). (**A**) t-SNE map visualization of embryos 10min - 3h old embryos with clusters identified by k-medoids clustering indicated by different colors. (**B**) Lineage analysis showing a single trajectory for all clusters (n > 1 embryos per cluster) leaving a total of 8 clusters (n = 84). (**C-D**) t-SNE map with coloring of individual dots according to the combined log2-transformed expression for 20 or 17 genes expressed during the minor (C) or major (D) wave of the ZGA (Supplemental_Table_S1.pdf). (**E-F**) Normalized read counts of minor (E) and major (F) ZGA genes for each embryo plotted along the pseudo-time order. The line represents the local regression of expression values on the ordered embryos. (**G-I**) Relative expression of select genes from manually staged embryos reported by (G) Sandler and Stathopoulos 2016 and (H) Lott *et al*. 2011 or (I) our computational age (pseudo-time). Gast, gastrulation. Gray background indicates the same developmental times included across datasets. For reference, the bar below the x-axis on I indicates clusters according to their color.

Plotting gene expression values along the pseudo-time axis provides a detailed insight into the dynamic expression patterns of these early transcribed genes. The published minor ZGA gene dataset (Kwasnieski et al. 2019) utilized in this analysis covered a tight developmental time window between NC 7 and 9, providing a static picture of an approximately 30 minutes long developmental window. In contrast, our analysis provides previously unprecedented resolution of the minor ZGA, showing a staggered onset of transcription for these genes (Supplemental_Fig_S2A,B.pdf). Intriguingly, many of the 20 minor ZGA genes share a similar sharp, transient peak of expression within the 3h time window, indicative of their role in early developmental processes. The exceptions are *E(spl)m4-BFM*, a member of the Notch signaling pathway, *sisA*, a gene involved in sex determination, and *CG6885*, a gene of unknown function. To identify the start of the major ZGA in our timeline, we used the combined expression levels of 17 genes that reportedly increase (≥5 fold) between NC 14A and NC 14B (Fig. 2D,F). These genes show increased expression at the transition from cluster 4 to 5 in our data (Fig. 2F).

Although the published gene list was curated using embryos within a 15 minute developmental time window, carefully staged according to time elapsed in interphase, nuclear elongation, and progression of cellularization (Sandler and Stathopoulos 2016), our continuous analysis shows that some of these genes actually increase transcription at unexpectedly early time points (Supplemental_Fig_S2C,D.pdf). Together, our results provide a detailed picture of the onset and dynamics of expression of previously reported ZGA transcript levels during early development.

To confirm the dynamic nature of expression patterns uncovered in our dataset, we compared the expression dynamics among a select group of genes that are known to be transcribed early (*screw, zerknullt, spitz, deadhead, stumps* and *yolkless*). These genes were previously shown to exhibit dynamic expression during development in different datasets that relied on the visual assessment and manual separation of samples into specific developmental categories or stages (Lott et al. 2011; Sandler and Stathopoulos 2016). We plotted the normalized expression levels for these genes, according to the stages disclosed in published datasets (Sandler (Fig. 2G) and Lott (Fig. 2H)) and according to our new computationally determined timeline (Fig. 2I). Graphs revealed that the transcriptional changes uncovered by our pseudo-time order are in good agreement with the previously published data. Pseudo-time order, cluster number and sample ID for each embryo are shown in Supplemental_Table_S2.pdf. Together, the results show that our method provides a high-resolution, time-sensitive picture of transcriptional events during early embryonic development.

### Single-cell RNA-sequencing reveals novel early transcribed genes

To determine if our method is able to identify novel early expressed genes, we compared expression of genes between embryos from cluster 1 and cluster 2 of the t-SNE map displayed in Fig. 2A. Within our dataset, we found the differential upregulation of transcription for 66 genes(padj<0.01, Log2FC>1) in this timeframe (Fig. 3A). Over-representation analysis (ORA) shows that these genes are involved in sex determination and developmental processes (Fig. 3B).

**Figure 3.**
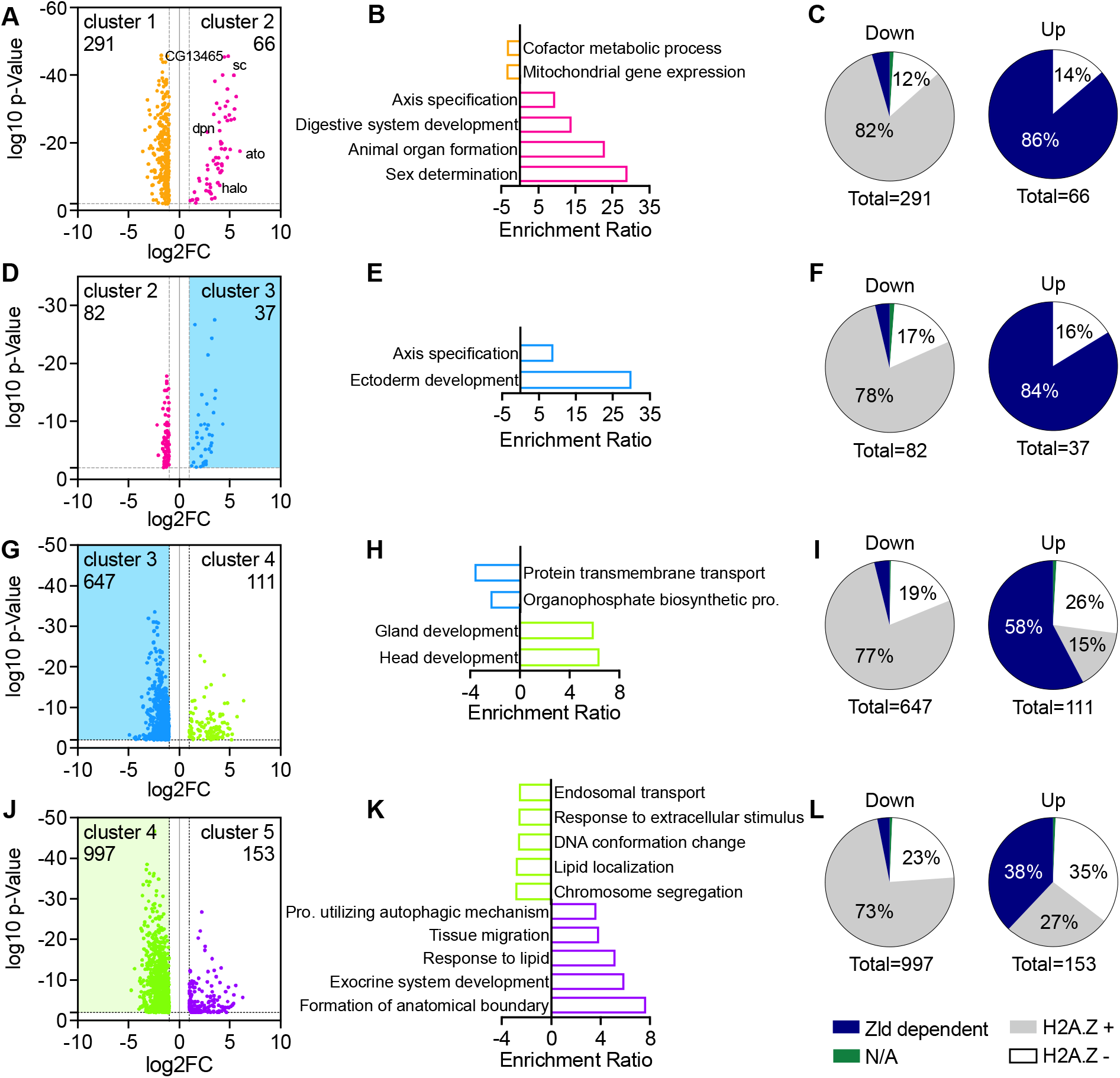
Differentially expressed genes, their related pathways and Zld or H2A.Z enrichment at TSS during the minor ZGA. (**A, D, G, J**) Volcano plots with significantly expressed genes (padj<0.01, Log2FC<-1 or >1) by comparing (A) cluster 1 versus 2, (D) cluster 2 versus 3, (G) cluster 3 versus 4, (J) cluster 4 versus 5 indicated in color. (A, D, G) The significantly changed unique transcripts not identified in previous cluster comparisons are represented by colored dots and the numbers are indicated in each volcano plot. (J) colored dots and number indicate all significantly expressed genes. (**B, E, H, K**) Significantly enriched pathways (FDR<0.05) by ORA on significantly expressed genes by comparing (B) cluster 1 versus 2, (E) cluster 2 versus 3, (H) cluster 3 versus 4, (K) cluster 4 versus 5. (**C, F, I, L**) Zld and H2A.Z enrichment at TSS (transcriptional start sites) of differentially expressed genes between (C) cluster 1 versus 2, (F) cluster 2 versus 3, (I) cluster 3 versus 4 or (L) cluster 4 versus 5. Zld data from Blythe and Wieschaus 2015 and H2A.Z enrichment from Ibarra-Morales *et al*. 2021. Genes not matching between datasets are shown as N/A. Pro., process.

To validate their early expression and determine the biological age at which these transcripts are activated, we compared our dataset to the most comprehensive dataset on early zygotic transcription published to date (Kwasnieski et al. 2019) and performed qPCR on a subset of genes in samples of hand staged fixed embryos spanning NC 6 to 11. The results from our single-embryo and qPCR analysis confirm the published evidence that *sc* is one of the earliest expressed genes at NC 7 (Supplemental_Fig_S3A.pdf). Our results also corroborate the early expression of all additional 19 genes previously reported to be expressed at NC 7-9. However, our analysis identified many other genes upregulated between cluster 1 and 2 which were previously reported to be expressed at significantly later timepoints - 31 genes at NC 9-10 and 15 genes at syncytial blastoderm. Importantly, qPCR results validated our single-embryo analysis approach and confirmed this earlier onset of expression for a randomly selected subset of genes (Supplemental_Fig_S3B,C,D,E.pdf; *ato* and *CG13465* were previously reported at NC 9-10 and *halo* and *dpn were* previously reported during syncytial blastoderm). We next wanted to exclude the possibility that these findings were due to the contamination of our qPCR samples with older embryos. To this end, we measured the levels of two gene transcripts (*hrg, bnb*) identified to be expressed at a later time point in our own temporal analysis (Fig. 3D; Supplemental_Fig_S3F,G.pdf, right panels) and in other published datasets (Lott et al. 2011; Kwasnieski et al. 2019). Using this approach, we detected no increase in expression for either *hrg* or *bnb* in the NC 7 and NC 8 samples (Supplemental_Fig_S3F,G.pdf, left panels), therefore, excluding the possibility of contamination of our NC 7 and 8 samples with older embryos. Further analysis revealed an additional upregulation of 37 genes between clusters 2 and 3, including *hrg* and *bnb* (Fig. 3D), and 111 genes between clusters 3 and 4 (Fig. 3G) with an enrichment in pathways related to early developmental processes (Fig. 3E,H). Our analysis identifies 214 genes that are significantly upregulated between clusters before the onset of the major ZGA. Taken together, the results show that our approach is able to identify the accurate onset of transcription of early expressed genes with high sensitivity.

We next compared expression between clusters 4 and 5 to identify genes activated at the beginning of the major ZGA (Fig. 3J). We identified 153 significantly upregulated transcripts in the cluster 5 embryos (padj<0.1, Log2FC>1). ORA revealed developmental pathways involved in tissue development, sex differentiation, and signaling pathways (Fig. 3K).

Previous studies have shown that a small subset of zygotically transcribed genes are dependent on the transcription factor Zelda (Zld) (Blythe and Wieschaus 2015), while a majority of zygotically transcribed genes are Zld independent but enriched for the histone variant H2A.Z (Ibarra-Morales et al. 2021).To explore Zld dependency and H2A.Z occupancy behavior over time in our dataset, we quantified the overlap between our up- and down-regulated transcripts in Fig. 3A,D,G,J in terms of being Zld dependent or independent while being bound or unbound by H2A.Z (Zld dependent, and Zld independent, which were divided into H2A.Z positive or negative) (Ibarra-Morales et al. 2021). Our analysis shows that the earliest minor ZGA genes are mostly Zld dependent (Fig. 3C,F) and that the share of Zelda dependent genes decreases sharply with the onset of the major ZGA. In contrast, the share of Zld independent genes, both H2A.Z positive and negative, increases with the onset of the major ZGA (Fig. 3I,L). These observations are not identified in down-regulated transcripts, which showed a distribution similar to all analyzed transcripts (Supplemental_Fig_S3H.pdf). In this way, we have demonstrated that our single-embryo RNA-seq methodology and analysis are a highly sensitive approach for identifying the accurate onset of gene transcription. Further, our analysis is able to define important transcriptional events and identify signatures of gene regulation during early development.

### mRNA decay of maternally deposited transcripts

In addition to transcription, our dataset also reveals patterns of maternal RNA decay. In order to identify maternally degraded mRNAs only, we compared cluster 1 (youngest embryos) with cluster 5 (onset of major ZGA) and selected only maternally deposited transcripts for analysis. Maternally deposited transcripts were defined as those with an averaged normalized read count >1 on the first 10 samples in our pseudo-time. Using this method, a total of 2621 significantly degraded transcripts were identified (padj<0.01, Log2FC<-1). 92% of our significantly degraded transcripts had also been shown to be degraded in a previously published dataset (classes II, III, IV, and V) (Fig. 4A) (Thomsen et al. 2010); only 35, 33, or 13 genes were within the Thomsen stable (class I), purely zygotic group of transcripts, or preloaded and transcribed, respectively. ORA of all 2621 degraded transcripts revealed mainly pathways related to metabolism (Supplemental_Fig_S4.pdf). This result likely reflects the elimination of transcripts important during oogenesis, but which are no longer needed for development. While patterns of transcript abundance differed before and after cluster 5, there is a clear inflection point at around value 50 of our pseudo-time. This time point coincides with the onset of the major ZGA. Maternal transcripts are deposited in oocytes at very different levels. To determine if degradation rates in the zygote are related to the initial quantity of deposited transcripts, we divided the significantly downregulated genes into 4 quartiles by their level of transcript abundance in cluster 1. We then determined the total number of normalized reads for each quartile in each individual embryo. Mean read counts plotted in Fig. 4B show the progressive nature of the maternal mRNA decay up to cluster 5. Plotting the ratio of cluster 5 to cluster 1 for the different quartiles (Fig. 4C) shows that the rate of decay is directly proportional to initial mRNA abundance, meaning that transcripts of low and high abundance are degraded at the same rate.

**Figure 4.**
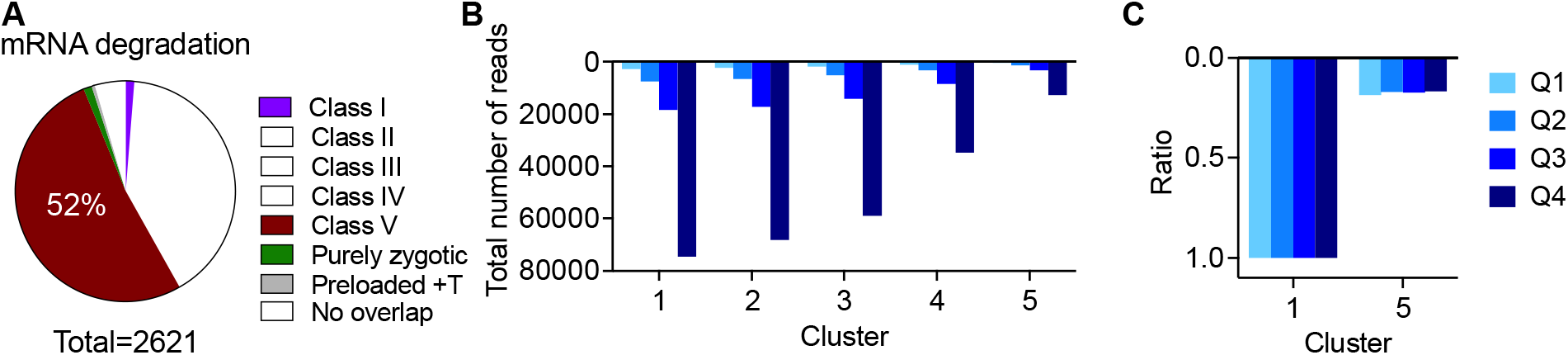
The continuous mRNA decay of maternally deposited transcripts. (**A**) Comparison of maternally deposited transcripts significantly decreased (padj<0.01, Log2FC<-1) by comparing cluster 1 versus 5 in our data and those previously reported by Thomsen *et al*., 2010. +T, and transcribed. (**B**) Mean read counts of all significantly decreased transcripts (n = 2621) by expression level group (Q, quartile) in each cluster. Q1, lowest 25%; Q4, highest 25% (**C**) Same data as in (B) showing the ratio of cluster 5 to cluster 1 by expression level.

Overall, we showed that maternal mRNA decay before the major ZGA is a progressive process. Degradation of maternal mRNA is proportional to transcript levels, suggesting that mRNA abundance is not related to degradation rate.

### X/Y Chromosome genotyping uncovers transcriptional dynamics of primary sex determination

Our prior analysis of the earliest transcribed genes indicates “Sex determination” as the most enriched pathway (Fig. 3B). The current model for primary sex determination is based on the tightly controlled sex specific expression levels of genes such as *Sex lethal* (*Sxl*) and *male-specific lethal* (*msl-2*) (Salz and Erickson 2010) during early development. This made us wonder if there are additional detectable differences in transcription between male and female embryos during our early developmental time window. To define the sex of each individual embryo, we isolated DNA from the organic phase after TRIzol™ extraction of RNA and performed qPCR using primers specific for the X- and Y-chromosome. Due to low DNA content of younger embryos, we get consistent PCR results only after embryo number 23 in our pseudo-time analysis. Based on these results, we categorized all embryos (after pseudo-time position 23) according to sex (Supplemental_Table_S2.pdf). To determine the differential expression of genes between male and female embryos, we used splineTimeR (see methods), which is particularly designed for identification of expression changes in longitudinal data. Our analysis identified 120 transcripts that were differentially expressed between male and female embryos (padj<0.01) (Fig. 5A). A large number of the differentially regulated genes are located on the X-chromosome (44%), whereas more than half the genes are located on autosomes and rDNA (56%). Several known regulators of primary sex determination such as *Sxl, sc, sisA*, and *msl-2* were also identified as significantly expressed in our analysis (Supplemental_Table_S3.pdf). Indeed, ORA shows that “Sex differentiation” is the most enriched pathway (Fig. 5B). We selected and plotted known regulators of sex determination using our pseudo-time scale (Fig. 5C,D). This analysis shows that differential transcription of *sc* and *sisA* (Fig. 5D) precedes the expression of *Sxl*. This agrees with the role of *sc* and *sisA* as activators of *Sxl* expression in females (Fig. 5C). Additionally, we identify other differentially expressed transcripts that precedes *Sxl* transcription, such as *CG14427*. Our differential expression analysis identified 21 transcripts upregulated in males and 99 in females (Supplemental_Table_S3.pdf). Examples for male specific expression are plotted in Fig. 5E. They show that the start of differential expression of *stonewall* (*stwl*, chromosome 3L), matches the start of differential expression of *Sxl* and *msl-2* and a pre-rRNA gene (*pre-rRNA:CR45847*) is expressed shortly after *Sxl* and *msl-2*, exclusively in males.

**Figure 5.**
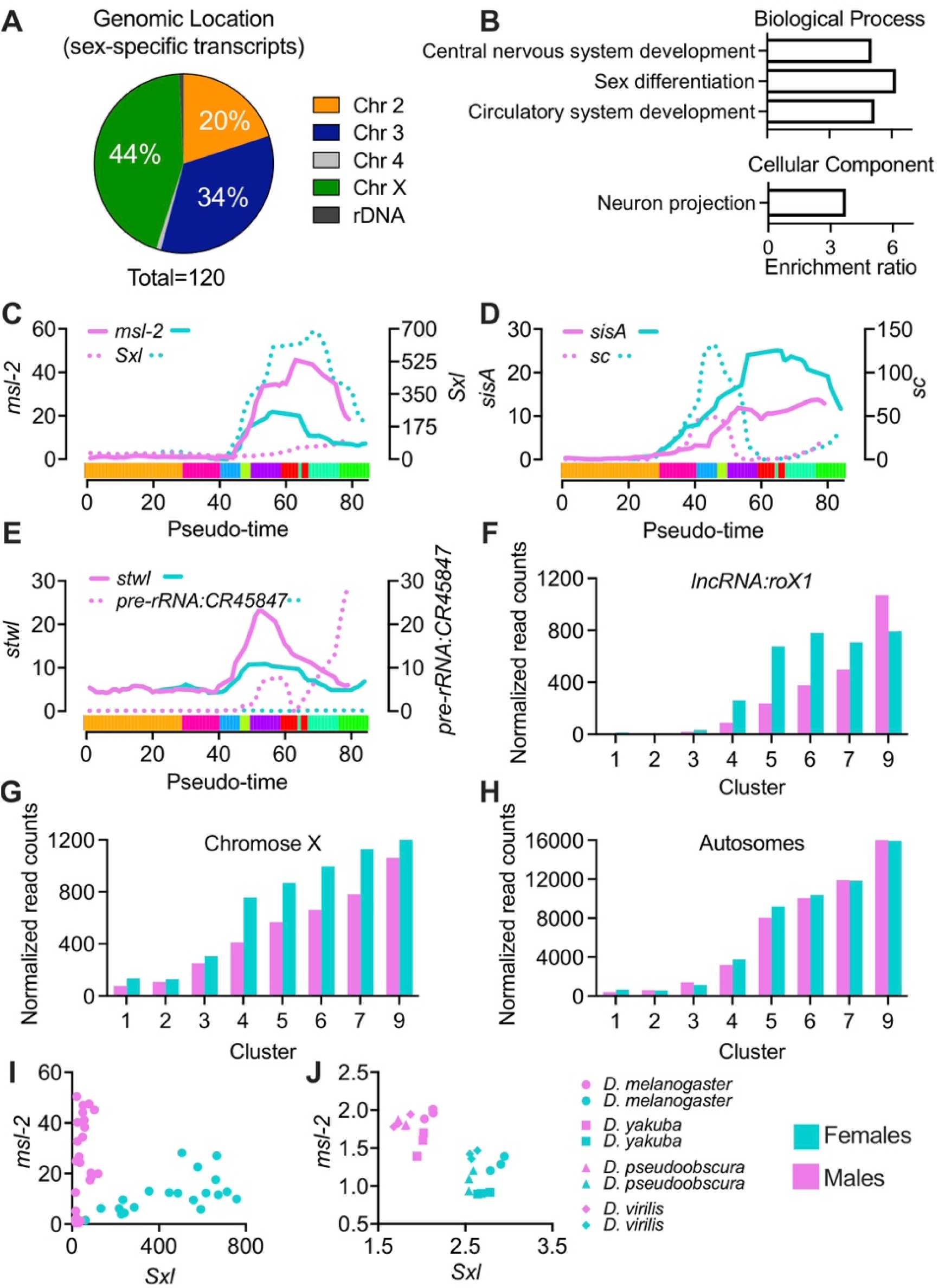
Sex-specific transcription and dosage compensation in the ∼3h embryo. (**A**) distribution of differentially expressed genes (padj<0.01) between male and female using splineTimeR according to their genomic location. (**B**) Significantly enriched pathways (FDR<0.05) of differentially expressed genes by ORA. (**C-E**) Smoothed normalized reads of selected transcripts. The colored bar along the x-axis shows clusters 1-9 from left to right, each in a different color for reference. (**F**) Average normalized reads for *lncRNA:roX1* or all zygotic transcripts (not maternally deposited) from male and female embryos within each cluster (**G**) for x-linked genes or (**H**) autosomal genes. (**I**) *msl-2* and *Sxl* normalized read counts of all male and female embryos in our data. (**J**) *msl-2* and *Sxl* FPKM (fragments per kilobase of transcript per million fragments mapped) of males and females from different *Drosophila* species reported in Paris *et al*., 2015.

Another important process linked to primary sex determination is dosage compensation, which assures equal expression of X-linked genes in males and females. In *Drosophila*, this is accomplished by the 2-fold upregulation of the X-chromosome in males. It has previously been reported that dosage compensation starts as early as NC 14C (Lott et al. 2011). To assess the onset of dosage compensation in our dataset, we excluded all maternally deposited transcripts and determined the total number of normalized reads for the remaining genes on the X-chromosome and autosomes for each individual embryo. Average read counts of male and female embryos within each cluster are plotted in Fig. 5G,H. From cluster 4 to 7, we observed a 1.5x higher expression of X-linked genes in female compared to male embryos, but no difference in autosomal reads. This difference was reduced to 1.1x in cluster 9 (gastrulation onset) (Fig. 5G), probably due to the start of canonical dosage compensation. Two components of the dosage compensation complex have been shown to have male-biased transcription, *msl-2* and *long non-coding RNA on the X 1* (*lncRNA:roX1*). *msl-2* is expressed at higher levels in males almost from the moment it is being transcribed (Fig. 5C) before we detect dosage compensation. *lncRNA:roX1* is higher in female embryos at first (Fig. 5F), which can be explained by its localization on the X-chromosome, levels only start to be higher in males once we see evidence for dosage compensation (cluster 8).

To further investigate how transcript levels are influenced by their chromosomal localization, we plotted early transcribed genes from our prior analysis (Fig. 2A,D) according to sex and chromosome location. Analysis shows that early transcribed genes from the X-chromosome, but not autosomes, tend to have higher levels in females compared with males (Supplemental_Fig_S5A,B,C.pdf). The time-point at which expression levels in females are higher than in males varies between genes, with some being detected as early as transcription of a gene starts (e.g. *ac, acheate*) and others occurring later in transcription (e.g. *run, runt*) (Supplemental_Fig_S5C.pdf).

Beyond the biological relevance of sex-specific transcription, we asked whether sex specific gene expression could serve as a tool to determine the sex of individual embryos. This approach would eliminate the need for the time-consuming genotyping approach. To this end, we plotted *Sxl* and *msl-2* transcript levels (Fig. 5I) and observed a clear separation of embryos according to their sex. We further validated this approach by applying it to a published single-embryo sequencing dataset (Paris et al. 2015), confirming that using *Sxl* and *msl-2* expression is sufficient to determine the sex in embryos of different *Drosophila* species (Fig. 5J). Of note, this approach only works in embryos after the onset of *Sxl* and *msl-2* transcription, around NC 12 and NC 14D (Supplemental_Fig_S5D.pdf).

Overall, our analysis detects sex-specific transcription as early as minor ZGA. Capitalizing on this differential gene expression, a simple strategy to sex embryos has been developed.

## DISCUSSION

Studying early development is challenging due to the rapid progression of biological processes and the limited amount of material available. To overcome these limitations, we developed a single-embryo RNA-seq and analysis approach, using the transcriptome as a measure of developmental progress (pseudo-time) to determine the biological age of the embryo. The high sensitivity of our method allows us to provide an accurate assessment of zygotic transcription and uncover the dynamic patterns for hundreds of genes. Our single-embryo approach also enables us to determine sex specific differences in transcript abundance. Utilizing these sex differences, we developed a new strategy to determine the sex of each embryo, without the need for genotyping. Together, we established an operationally simple method to document gene expression changes at unprecedented resolution and provide a continuous assessment of transcriptional processes during early development.

### An operationally simple, single-embryo sequencing method

Previous studies investigating zygotic transcription relied on precise collection time windows and/or elaborate manual staging of the embryos under a microscope. *Drosophila* females, however, are known to lay unfertilized eggs and withhold embryos (Markow et al. 2009). Human error in staging of embryos and irregular laying patterns of females can lead to the inclusion of mis-staged embryos in the analysis. Recent studies have highlighted the advantages of single embryo sequencing approaches over working with pooled samples (Paris et al. 2015; Liu et al. 2020), specifically the ability to detect and exclude older embryos from the analysis, therefore providing more accurate data. In this work, we present an optimized single-cell sequencing protocol for use with *Drosophila* embryos and a single-cell bioinformatic pipeline analysis. We assign a computationally derived age to each embryo, based upon their transcriptome, thereby circumventing the need for elaborate and error prone staging procedures. Indeed, we show that our computationally derived pseudo-time reflects the biological age of the embryos, by comparison with previously established datasets. In addition, our protocol reduces reagent and sequencing costs due to the low volume nature of the experiments and the inclusion of UMIs (Sagar et al. 2018). Further, it requires no special instrumentation, beyond a micro-pipetting device, and the analysis utilizes established tools (Herman et al. 2018). Together these advantages make the method reported here the most accessible methodology developed to date, opening up this type of research to almost any *Drosophila* lab. This single embryo sequencing approach, ultimately, will lead to an improved reproducibility of developmental studies between experiments and laboratories.

### An accurate characterization of early transcriptional events

Our pseudo-time approach allows us to identify the exact onset of transcription even for lowly expressed genes. Our results reveal that previously reported, as well as many novel transcripts, are expressed as early as NC7. One example is the early expression of *halo*, a cofactor for the molecular motor, kinesin and a regulator of lipid droplet movement, which was previously reported to be actively transcribed during syncytial blastoderm (after NC 11) (Kwasnieski et al. 2019), but was identified as one of the earliest transcribed genes in our dataset. Our analysis also reveals the dynamic nature of transcriptional events. It provides information about expression for thousands of genes at a temporal resolution unchallenged by other methods. Recently, a single-cell dataset was published covering all of embryogenesis and providing insights into cell type specific transcriptional changes during development (Calderon et al. 2022). While this dataset provides an incredibly detailed insight into *Drosophila* development, it only detected a median of 399 UMIs and 274 genes per cell, likely only covering very highly expressed genes. In contrast, we detected a median of over 600,000 UMIs and identified over 7200 genes per embryo, leading to a total 9777 identified genes across the whole dataset. Thus, our dataset gives a much more complete picture of transcriptional changes during early development. While our dataset lacks the cell type specific information, this might be of minor relevance during early development. It is known that nuclei share the same cytoplasm until cellularization, and the single-cell sequencing dataset only identified 3 different types of cells (anlage in statu nascendi, aminoesra anlage, ectoderm anlage) during the first 4h of development. That said, we acknowledge that cell specific expression patterns become more important at later stages in development, and while our method does not allow for this kind of analysis, it is much more accessible than single-cell sequencing of thousands of cells at different developmental time points.

### Sex-specific gene expression

Our single-embryo method also allows us to distinguish between male and female embryos, opening the possibility of investigations into sex-specific transcriptional effects. X-signal elements (XSEs) have been shown to control the early sex specific expression of *Sxl* and to drive primary sex determination. We show differential expression of *sc* and *sisA*, two strong XSEs, from the very first moment of ZGA (NC 7). Of note, CG14427, an X-linked gene with unknown function, is also differentially expressed between males and females starting at NC 7, making it a potential candidate as a novel XSE. Our data also allows for further insights into primary sex determination. An early expressed XSE, *runt*, was previously reported to undergo a non-canonical form of dosage compensation and expression which was proposed to be under the direct control of *Sxl* (Gergen 1987; Smith et al. 2001). Our results show that *runt* is one of the earliest transcribed genes at similar levels in males and females, preceding *Sxl* transcription. This argues against the role of *Sxl* in controlling *runt* expression. However, our data also show a differential expression of *runt* between male and females after *Sxl* peak expression. As such, it is possible that rather than controlling overall *runt* expression, *Sxl* only controls differential expression of *runt* later in development. Surprisingly, we also identified several autosomal encoded genes as differentially expressed between males and females. While differential expression of X-chromosomal genes in females can be explained by their different dosage (2X in females vs 1X in males), this is not the case for autosomal genes, which are present at equal dosage in both males and females. These results suggest additional players in primary sex determination. Further studies will be needed to confirm these results and investigate the underlying mechanisms. Importantly, our newly developed strategy to determine the sex of single-embryos by using the expression of known regulators of primary sex determination (*Sxl* and *msl-2)* eliminates the need for elaborate genotyping procedures in future sequencing datasets.

Taken together, we believe our method is the most accessible, high-throughput, transcriptomic technology published to date to study early gene expression in *Drosophila*. We suggest that our methodology provides the optimal tool to investigate the transcriptional consequences of mutations in developmental genes, providing gene expression data at unprecedented depth and temporal resolution.

## METHODS

### Fly stocks and embryo collection

Drosophila genetic reference panel (DGRP) 737 line from Bloomington Stock Center (#83729) was kept in incubators at 25 °C with 60% humidity and a 12-hour light-dark cycle. All flies were raised at constant densities on standardized cornmeal food (Bloomington recipe), Fly food M (LabExpress, Michigan, USA), and transferred into cages 1-2 days after eclosion. Food plates were changed and discarded twice before embryo collection started on 8-9 day old flies. DGRP_737 line showed minimal egg laying (n = 0-2) in the first 30 minutes (data not shown), therefore, plates were changed every 90 minutes and processed immediately (0-1 h embryos) or incubated 1 or 2 more hours at 25 °C (1-2 h or 2-3 h embryos, respectively). Embryos were transferred into a pluriStrainer® 150 µM cell strainer (pluriSelect, USA) and washed with tap water, dechorionated by incubation in 3% sodium hypochlorite (PURE BRIGHT® bleach, KIK international LLC) for 4 min, washed in 120 mM NaCl (Sigma-Aldrich, USA), 0.03% Triton X-100 (Fisher Scientific, USA) solution, and finally washed in ultrapure water (PURELAB® Ultra, ELGA). For RNA-sequencing (RNA-seq), single embryos were transferred into 2 ml screw-cap microtubes using a 20/0 liner brush (Royal & Langnickel®, USA), snap-frozen on dry ice, and stored at -80 °C. For embryo fixation, samples were processed immediately.

### RNA isolation

Single embryos were homogenized in 500 µL TRIzol™ (Invitrogen, USA) by bead-beating with 0.2 g lysing matrix D beads (MP Biomedicals, USA) at 6 m/s for 30 seconds using the FastPrep-24™ instrument (MP Biomedicals, USA). RNA was then isolated following a miniaturized version of the manufacturer’s instructions. Briefly, 100 µL chloroform (Sigma-Aldrich, USA) was added, samples mixed by vortex, incubated 2 min at room temperature (RT), and centrifuged for 15 minutes at 12,000 × g at 4 °C to recover RNA-containing aqueous phase in a fresh 1.5 ml microtube. At this step, the organic phase was stored at -80 °C for later DNA extraction. RNA was precipitated by adding 250 µL ice-cold isopropanol (Sigma-Aldrich, USA) and 2 µL GlycoBlue™ (Thermo Fisher, USA), samples mixed by hand, incubated for 10 min at RT, and centrifuged for 10 minutes at 12,000 × g at 4°C. RNA pellets were washed with 1 ml 75% (v/v) ethanol (Pharmco, USA), dried and stored at -80 °C until further use. This same protocol was followed to isolate RNA from fixed embryos using 1 ml TRIzol™ and proportional changes in chloroform and isopropanol.

### Library preparation and RNA-seq

RNA-seq was carried out following a miniaturized version of the sensitive highly-multiplexed single-cell RNA-seq (CEL-Seq2) protocol (Hashimshony et al. 2016) using the I.DOT liquid handler (CELLINK). Dried RNA was resuspended in 8 µL nuclease-free water (Invitrogen, USA) and 120 nL dispensed into a 384-well plate holding 240 nL of primer-mix including 192 different cell barcodes with unique molecular identifiers (UMI). Subsequent steps and reagents are detailed in Sagar *et al*., 2018, except that libraries were diluted 1:10 (cDNA:H_2_O) before an 11-cycle amplification. Paired-end sequencing (150 bp) was performed using the NovaSeq 6000 instrument (Illumina) by the Genomics Core at Van Andel Institute. Sequencing depth in each single embryo was between 6.4-6.8 M reads that passed quality control, with 96% of the sequences with a quality score ≥ 30 (FastQC version 0.11.9) (Andrew 2010).

### RNA-seq data analysis and functional enrichment

RNA-seq read counts were demultiplexed, mapped to the Berkeley Drosophila Genome Project assembly release 6.28 (Ensembl release 100) reference genome (Hoskins et al. 2015), UMI-deduplicated, and counted using STAR 2.7.8a (mode STARsolo). Gene symbols were updated using release FB2022_04. Samples with a total transcript read count < 250,000 or transcripts with < 3 read counts on < 5 samples were filtered out from the analysis. Read count normalization, computation of a distance matrix, sample clustering, transcriptome entropy calculi, generation of a lineage tree, and pseudo-temporal order of samples was carried out using R packages RaceID version 0.2.6 and FateID version 0.2.2 (Herman et al. 2018). Raw expression values of unsupervised clusters were compared by the RaceID3 internal approach akin to DESeq2. Transcripts with a Benjamini-Hochberg adjusted p-value (padj) < 0.01 and a log2 fold-change (Log2FC) <-1 or >1 were considered to be differentially expressed. The source code for this analysis can be found in Supplemental_Table_S4.pdf. All functional enrichment analysis were carried out by over-representation analysis (ORA) using the WEB-based GEne SeT AnaLysis Toolkit (WebGestalt) (Liao et al. 2019). To simplify results, redundancy reduction by affinity propagation was applied in every analysis. Only results with a false discovery rate (FDR) ≤ 0.5 are shown.

### Fixation, staining and staging of embryos

Dechlorinated embryos were transferred to a 1.5 ml microtube and mixed in 362.5 µL PBT (0.3% Triton X-100 in Gibco™ 1x phosphate-buffered saline (PBS), pH 7.4), 12.5 µL 10x PBS, and 125 µL 16% formaldehyde, methanol-free (w/v) (Thermo Fisher Scientific, USA). Embryos in the 4% formaldehyde fixing solution (w/v) were shaken for 15 min at 200 rpm using a mini rotator/shaker (Thermo Scientific). Fixing solution was discarded, 500 µL heptane (Sigma-Aldrich, USA) and 500 µL methanol (Fisher Scientific, USA) added, and samples vigorously shaken by hand/vortex for 2 min. Heptane, methanol and embryos in the interphase were removed and discarded. Samples were washed 3 times with methanol before resuspension in 1 ml PBT containing 1 µL Hoeschst 33342 (Thermo Scientific, USA). After a 10 min incubation at RT, 2 × 1 min and 1 × 10 min washes with 1 ml PTB were carried out to remove excess dye. Embryos were then staged using the ECLIPSE Ts2 microscope (Nikon) based on Foe *et al*., 1993, nuclear cycle divisions and images from others (Jiménez-Guri et al. 2014; Kotadia et al. 2010). Embryos in PBT were kept on ice during staging. Finally, PBT was removed, TRIzol™ added to pooled embryos and samples stored at - 80 °C until RNA isolation.

### Reverse transcription

Dried RNA from stage embryos was resuspended in 9 µL nuclease-free water and 1 µL used for quantification by NanoDrop™ One/OneC spectrophotometer (Thermo Scientific). The remaining RNA (< 3 µg) was treated with 2 U TURBO™ DNase (Invitrogen, USA) following manufacturer’s instructions. RNA was then incubated at 70 °C with 1.5 µg oligo(dT)_12-18_ (Invitrogen, USA) and immediately chilled on ice. Reverse transcription was carried out using moloney-murine leukemia virus (M-MLV) reverse transcriptase kit (Promega, USA). Reverse transcription was completed in a 30 µL final volume reaction containing 400 U M-MLV and 1 mM dNTP mix (Invitrogen, USA) after serial incubations at 40 °C for 60 min and 90 °C for 10 min. cDNA was chilled on ice and diluted to a concentration of 20 ng/µL (1 µg input RNA/50 µL).

### DNA extraction

DNA extraction was performed with a modified version of the manufacturer’s instructions (TRIzol™, Invitrogen, USA). The frozen organic phase of each embryo after RNA extraction was thawed at RT for 3 min and transferred to a fresh 1.5 ml microtube to remove beads from samples. 2 µL GlycoBlue™ were added, samples mixed by inverting tube 5 times, 150 µL 100% ethanol (Pharmco, USA) were added, and samples mixed again. After a 3 min incubation at RT samples were centrifuged 5 min at 7,000 g at 4 °C and the phenol-ethanol supernatant discarded. DNA pellets were washed in 500 µL 0.1 M sodium citrate in 10% ethanol and incubated 30 min mixing every 10 min. Samples were centrifuged for 5 min at 7,000 g at 4 °C and supernatant discarded. Wash with 0.1 M sodium citrate was repeated once, and pellets resuspended in 1 ml 75% ethanol. Then, 2 µL GlycoBlue™ were added and samples incubated for 10 min mixing every 2-5 min. Samples were centrifuged 5 min at 7,000 g at 4 °C, supernatant was discarded and pellet air dried before resuspension in 20 µL 8 mM NaOh in H2O (w/v). DNA samples were incubated at 80 °C for 10 min mixing every 2 min by vortex, chilled immediately on ice for 5 min, and stored at 80 °C.

### qPCR and data analysis

qPCR was carried out in a 20 µL final volume reaction using SsoAdvanced Universal SYBR Green Supermix (Bio-Rad, USA), bespoke forward/reverse primers (0.3 μM/each) (Supplemental_Table_S5.pdf) and 2 µL DNA (1/10 embryo) or 160 ng/μl cDNA. Pre-incubation at 98 °C for 3 min for DNA or 30 s for cDNA, 45 cycle amplification, and melting curve were performed using CFX96 touch real-time PCR detection system (Bio-Rad). Each amplification cycle included denaturation at 95 °C for 10 s and a combined annealing/extension at 60 °C for 30 s. Specificity of qPCR reactions was assessed by the presence of a single peak in the melting curve, which was generated by acquiring fluorescence data every 0.5 °C change in temperature from 65 °C to 95 °C. All qPCR reactions were performed in duplicate (technical replication). DNA samples that amplified the X and Y-chromosome in both duplicates at similar cycle threshold values were categorized as males. DNA samples that amplified the X but not the Y-chromosome were categorized as females. For cDNA samples, mRNA expression in each duplicate was calculated using the cycle threshold values by the standard curve method (Cikos et al. 2007). Expression was then divided by the square root of CG6707 (FBgn0036058) multiplied by Pgam5 (FBgn0023517), two transcripts with the lowest variability until around NC 14D in our RNA-seq data.

### Sex-specific transcription

RNA-seq normalized read counts of each transcript were compared between male and female embryos using splineTimeR version 1.24.0 (Michna et al. 2016). Every embryo was considered a replicate in every cluster (timepoints). Transcripts with a Benjamini-Hochberg adjusted p-value (padj) < 0.01 were considered significantly expressed. The source code for this analysis can be found in Supplemental_Table_S4.pdf. Due to the split of the pseudo-time into male and females, normalized reads were smoothed by averaging 5 neighboring samples and a second order of the smoothing polynomial using Prism 9 version 9.4.1.

## DATA ACCESS

Raw sequencing files and processed data files from this study have been deposited at NCBI Gene Expression Omnibus (GEO) under accession number GSE214118 and will be released after publication in a peer reviewed journal. Normalized reads, gene details, and metadata for sex-specific analysis can be found in Supplemental_Table_S6.xlsx.

## ACKNOWLEDGMENTS

We thank Carolyn Anderson for critical reading and editorial feedback on the manuscript. We are grateful to all members of the Lempradl lab for helpful discussions. We thank the Van Andel Institute Genomics Core, especially Marc Wegener and Marie Adams, for their assistance with RNA-sequencing. We thank the VAI Bioinformatics and Biostatistics Core for supporting the analyses. We thank J. Andrew Pospisilik for supporting the project.

## AUTHOR CONTRIBUTIONS

J.E.P-M. and A.L designed and directed the study. J.E.P-M., J.W, L.E performed experiments. J.E.P-M., K.L. and A.L .analyzed the data. J.E.P-M. and A.L. wrote the manuscript.

