## Supplemental Material for "Continuous transcriptome analysis reveals novel patterns of early gene expression in *Drosophila* embryos"

Pérez-Mojica *et al*.

This PDF file includes the following:

Other supplemental material for this manuscript includes:

Supplemental_Table_S6.xlsx (Normalized read counts by pseudo-time order, genomic location of each gene and metadata for sex-specific transcription analysis)

Supplemental Figure S1

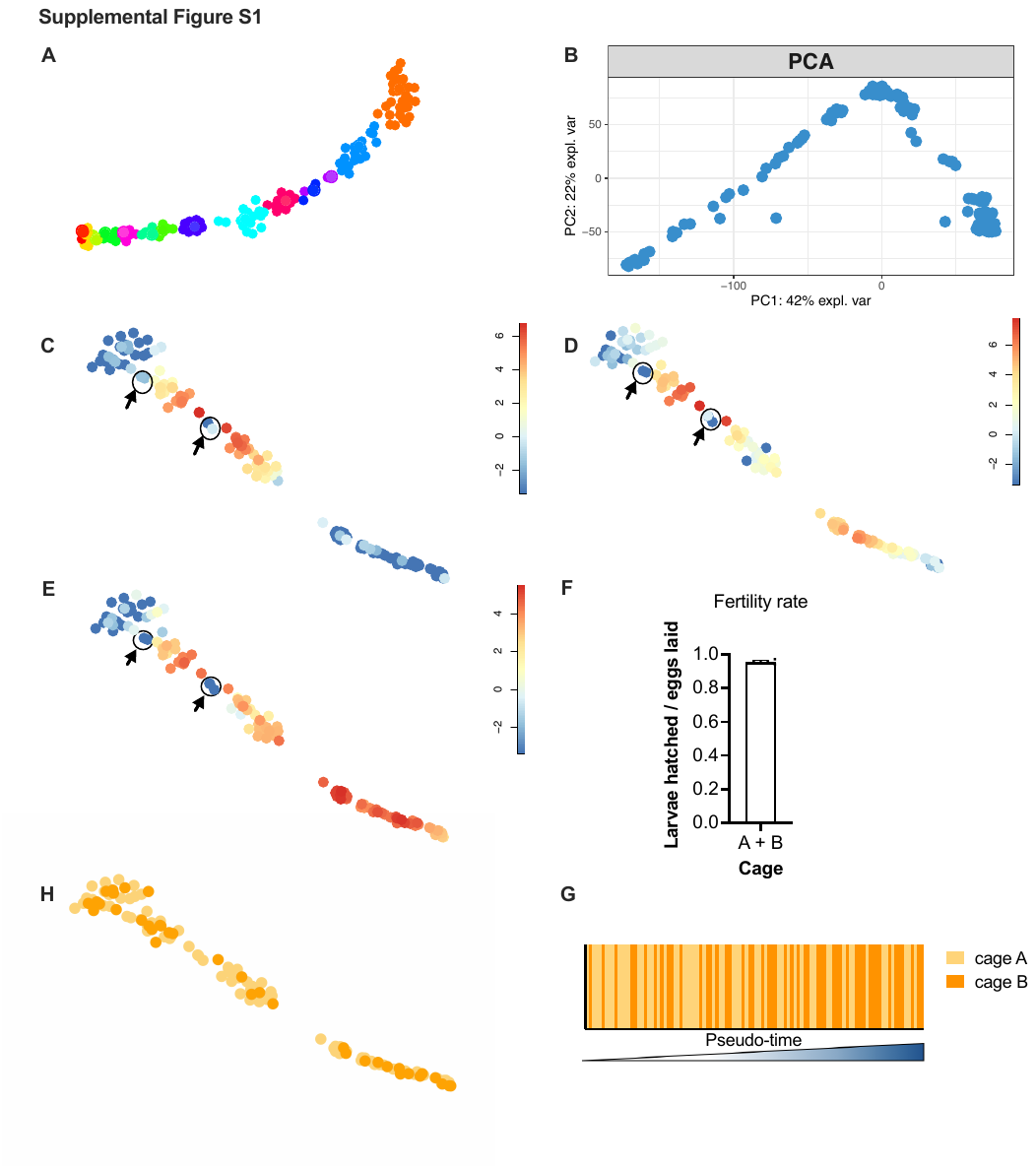

**Supplemental Figure S1.**

**Alternative dimensionality reductions, identification of unfertilized eggs and analysis of batch effects in RNA-seq data.** (**A**) Fruchterman-Reingold layout or (**B**) PCA dimensionality reduction of raw RNA-seq data. (**C-E**) t-SNE maps of log2-transformed expression values with arrows and circles indicating excluded samples (n = 5) due to low levels of (**C**) *screw* (*scw*), (**D**) *scute* (*sc*) and (**E**) *escargot* (*esg*) compared to adjacent embryos on the t-SNE map and pseudo-time. (**F**) The ratio of larvae hatched from embryos taken from the two cages (A and B) used for RNA-seq on the same day of collection averaged 0.95 (n= 6, 3 replicates of ~150 embryos per cage). (**H**) t-SNE map or (**G**) pseudo-time showing cage origin (A or B) of each sample.

Supplemental Figure S2

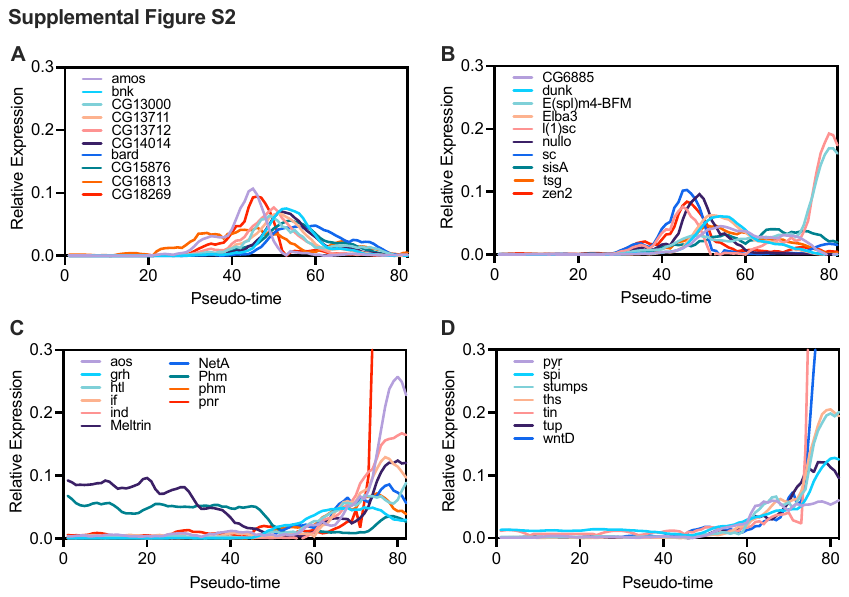

**Supplemental Figure S2.**

**The continuous sequence of the ZGA, a detailed look.** Graphs show the loess-smoothed expression on reads normalized to sum 1 for (**A-B**) 20 genes reported to start transcription during NC 7-9 by Lott, *et al.*, 2011 or (**C-D**) 17 genes reported to increase >5-fold expression from NC 14A to NC 14B by Sandler and Stathopoulos, 2016.

Supplemental Figure S3

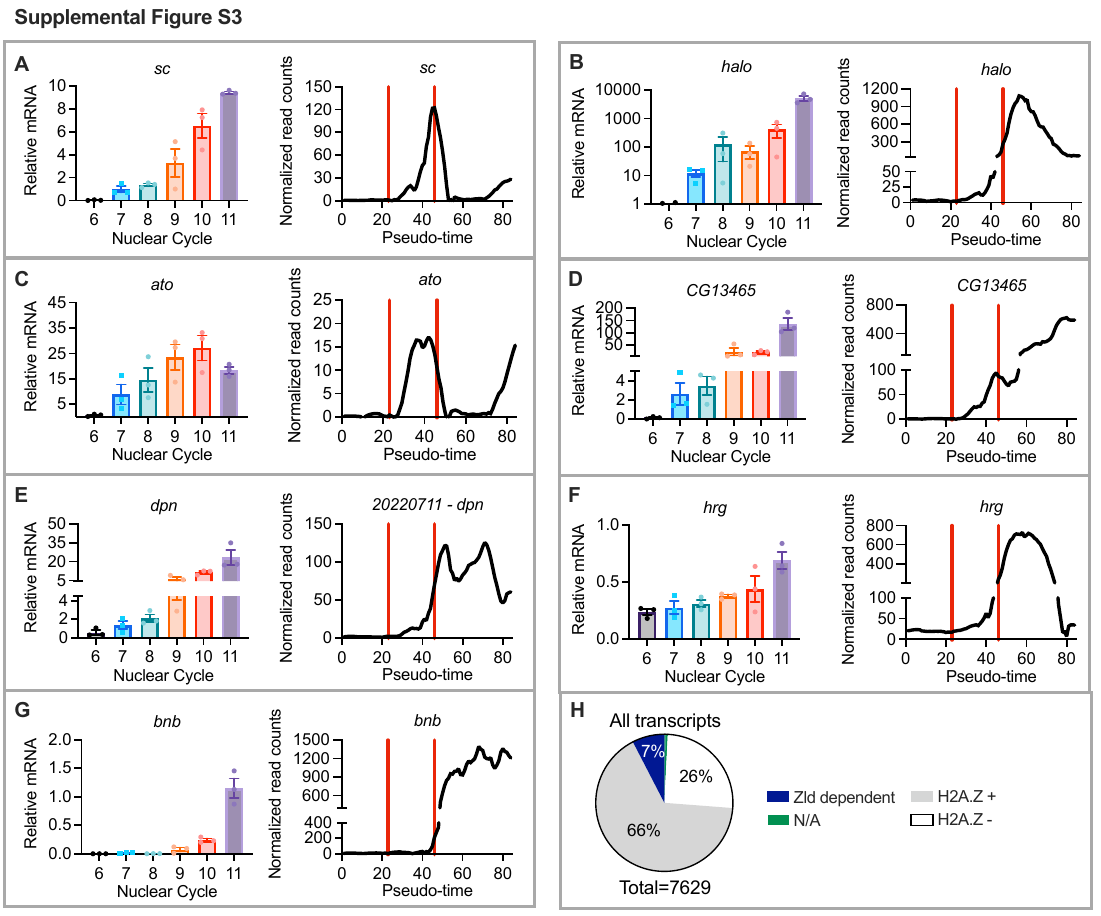

**Supplemental Figure S3.**

**qPCR results for selected genes and genome distribution of Zelda or H2A.Z.** (**A-G**) On the left panel, qPCR from fixed and hand-staged embryos at NC 6-11. On the right panel, smoothed normalized reads in our pseudo-time. Normalized reads were smoothed by averaging 5 neighboring samples and a second order of the smoothing polynomial using Prism 9 version 9.4.1. Vertical red lines indicate an approximate window covering NC 6-11 in our pseudo-time. Gene symbol and gene name: *sc*, *scute*; *halo*, *halo*; *ato*, *atonal*; *CG13465*, no name; *dpn*, *deadpan*; *hrg*, *hiiragi*; *bnb*, *bangles and beads*. (**H**) Genome distribution of Zld or H2A.Z in all transcripts detected up to ~3h embryos. Zelda bounding data from Blythe and Wieschaus, 2015 and H2A.Z enrichment from Ibarra-Morales *et al*. 2021. Genes not matching between datasets are shown as N/A.

Supplemental Figure S4

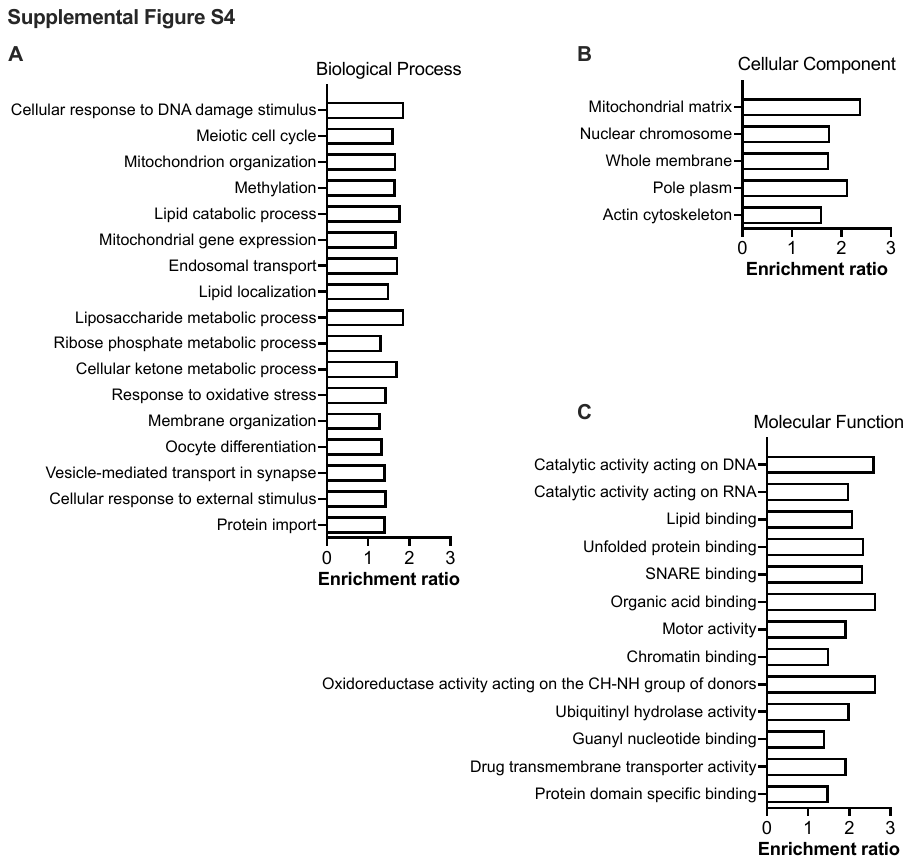

**Supplemental Figure 4.**

**Pathway analysis of maternally deposited mRNAs degraded upon major ZGA.** (**A-C**) ORA on all (n = 262) significantly decreased genes by comparing cluster 1 versus 5 (padj<0.01, Log2FC<-1)

Supplemental Figure S5

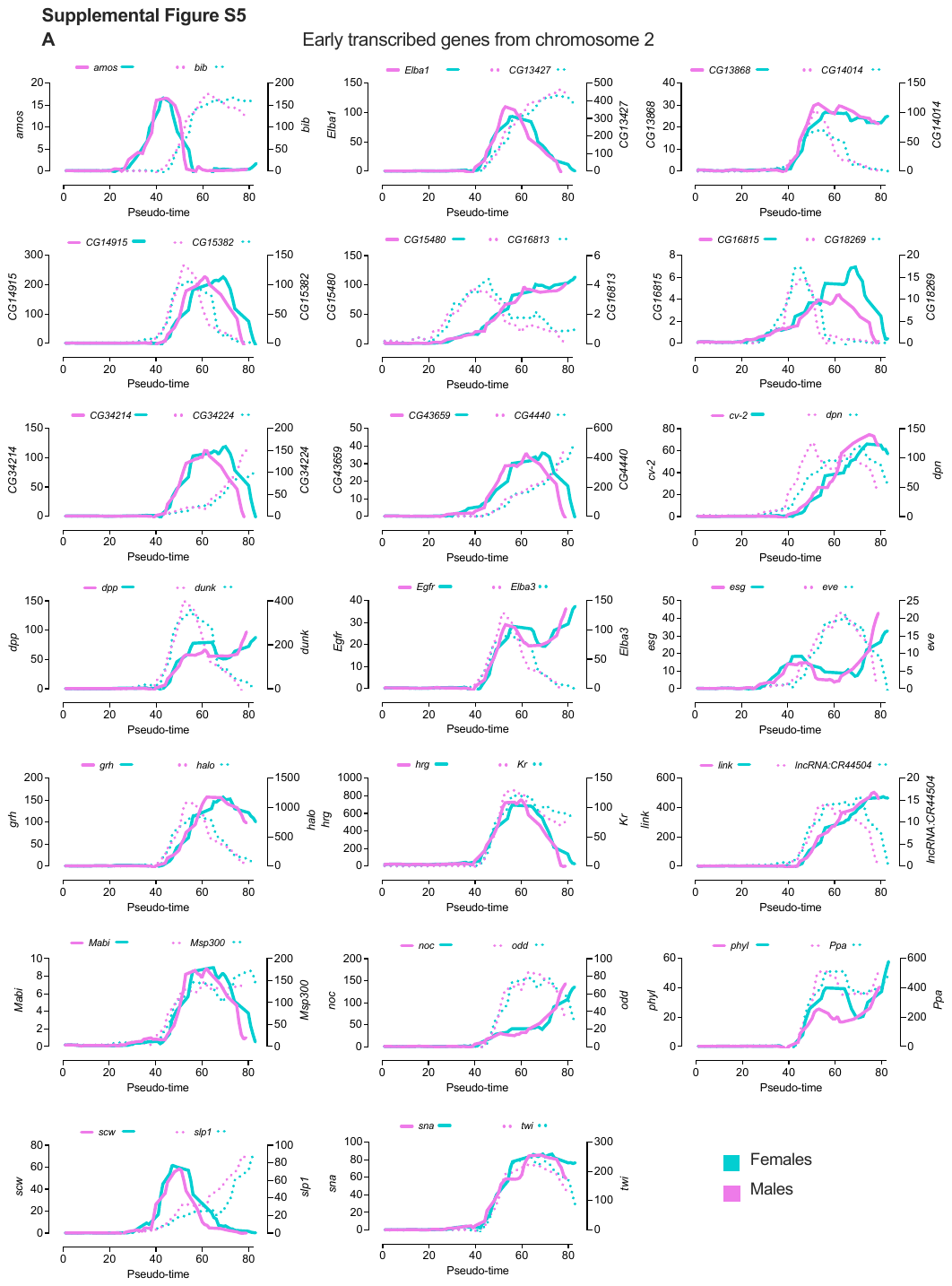

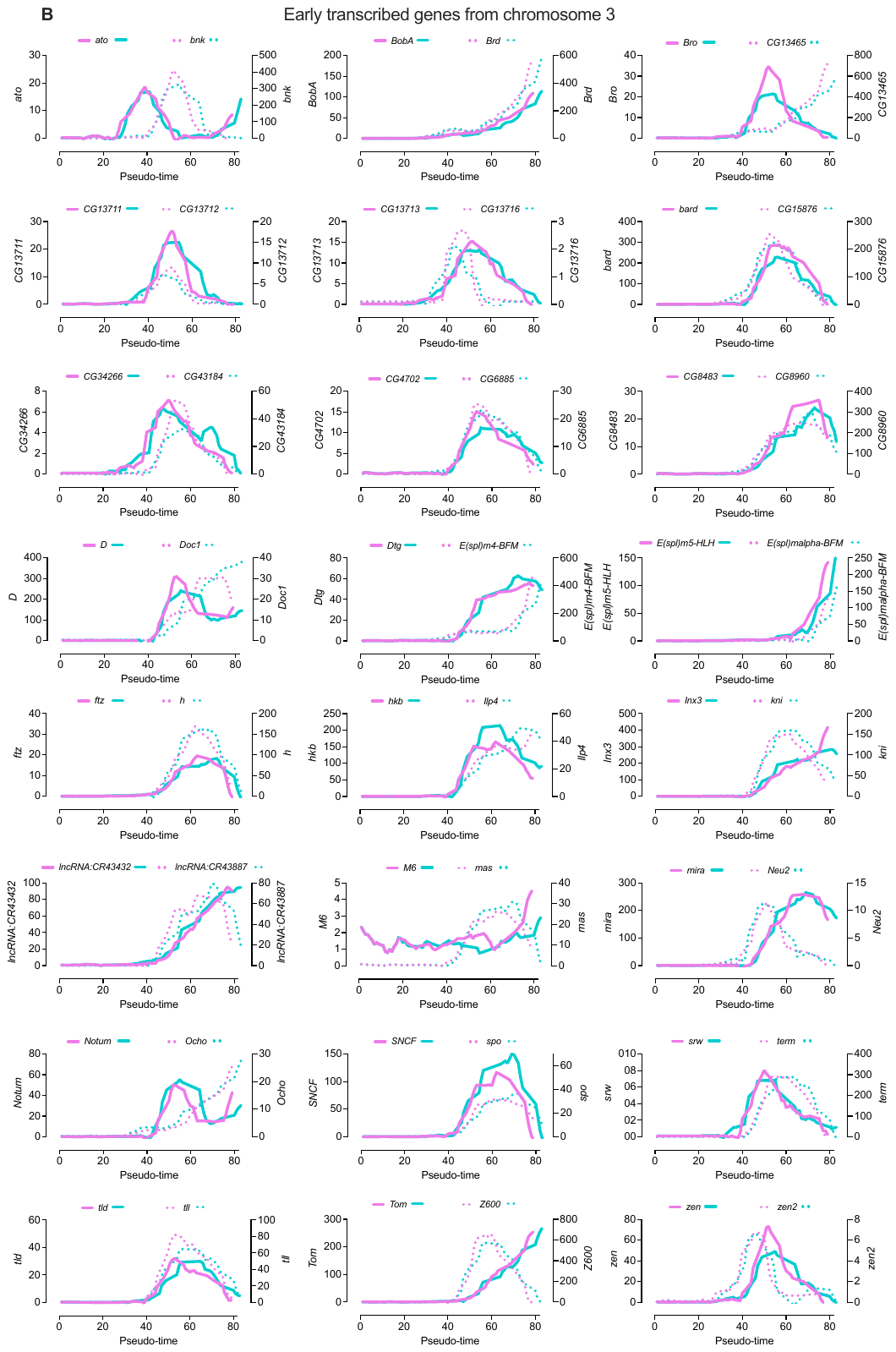

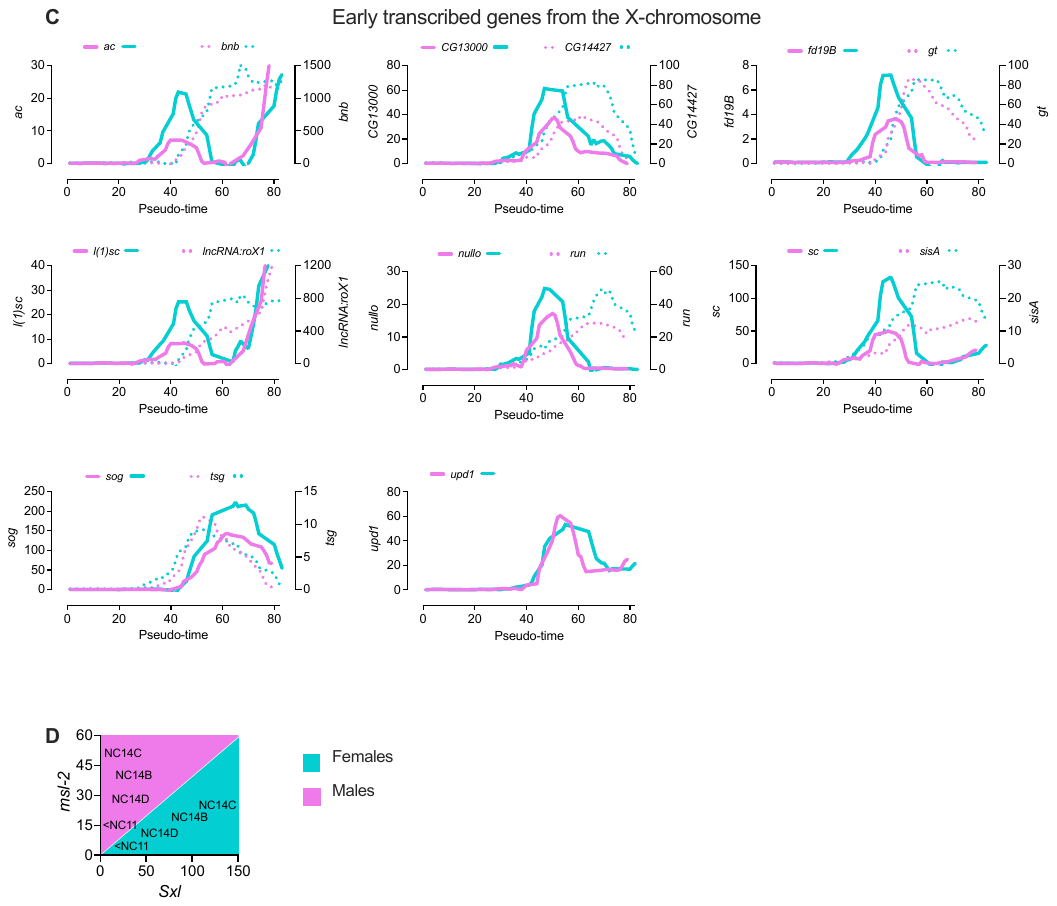

**Supplemental Figure S5**

**Sex-specific transcription in early development and developed strategy to sex embryos.** (**A-C**) Smoothed normalized reads of significantly increased transcripts (padj<0.01, Log2FC>1) by comparing cluster 1 versus 2 and cluster 2 versus 3. Genes were grouped according to their genomic localization on (A) chromosome 2, (B) chromosome 3, or (C) the X-chromosome. (**D**) Cartoon showing how males and female embryos cluster by plotting *Sxl* expression on the x-axis and *msl-2* expression on the y-axis.

Supplemental Table S1.

Previously reported genes during the minor and major ZGA.

| Minor ZGA (NC 7-9) | | |  | Major ZGA (NC 14A-14B) | | |
| --- | --- | --- | --- | --- | --- | --- |
| FlybaseID |  | Gene symbol |  | FlybaseID |  | Gene symbol |
| FBgn0002629 |  | E(spl)m4-BFM |  | FBgn0020299 |  | stumps |
| FBgn0031621 |  | Elba3 |  | FBgn0003963 |  | ush |
| FBgn0002561 |  | l(1)sc |  | FBgn0265140 |  | Meltrin |
| FBgn0004389 |  | bnk |  | FBgn0001250 |  | if |
| FBgn0003270 |  | amos |  | FBgn0025776 |  | ind |
| FBgn0083973 |  | dunk |  | FBgn0004569 |  | aos |
| FBgn0004143 |  | nullo |  | FBgn0033652 |  | ths |
| FBgn0003411 |  | sisA |  | FBgn0003117 |  | pnr |
| FBgn0004170 |  | sc |  | FBgn0005672 |  | spi |
| FBgn0003865 |  | tsg |  | FBgn0033649 |  | pyr |
| FBgn0004054 |  | zen2 |  | FBgn0259211 |  | grh |
| FBgn0030807 |  | CG13000 |  | FBgn0004959 |  | phm |
| FBgn0035572 |  | CG13711 |  | FBgn0038134 |  | wntD |
| FBgn0035570 |  | CG13712 |  | FBgn0004110 |  | tin |
| FBgn0031718 |  | CG14014 |  | FBgn0010389 |  | htl |
| FBgn0038566 |  | bard |  | FBgn0003896 |  | tup |
| FBgn0035569 |  | CG15876 |  | FBgn0015773 |  | NetA |
| FBgn0031719 |  | CG18269 |  | N/A |  | N/A |
| FBgn0036810 |  | CG6885 |  | N/A |  | N/A |
| FBgn0032490 |  | CG16813 |  | N/A |  | N/A |
| Genes transcribed during the minor ZGA were taken from Kwasnieski et al. 2019. Genes transcribed during the major ZGA were taken from Sandler and Stathopoulos 2016. | | | | | | |

Supplemental Table S2.

Pseudo-time order, cluster number, sample ID and sex for each embryo.

| Pseudo-time order | Cluster | Sample ID | Sex |  | Pseudo-time order | Cluster | Sample ID | Sex |
| --- | --- | --- | --- | --- | --- | --- | --- | --- |
| 1 | 1 | X1A.22 | NO DNA |  | 43 | 3 | X2A.04 | FEMALE |
| 2 | 1 | X1B.04 | NO DNA |  | 44 | 3 | X2A.32 | FEMALE |
| 3 | 1 | X2A.07 | NO DNA |  | 45 | 3 | X2A.30 | MALE |
| 4 | 1 | X1B.20 | NO DNA |  | 46 | 3 | X2A.15 | MALE |
| 5 | 1 | X1A.19 | NO DNA |  | 47 | 3 | X2B.20 | FEMALE |
| 6 | 1 | X1A.02 | NO DNA |  | 48 | 4 | X2A.21 | FEMALE |
| 7 | 1 | X1A.20 | NO DNA |  | 49 | 4 | X2A.18 | MALE |
| 8 | 1 | X1A.25 | NO DNA |  | 50 | 4 | X2B.24 | FEMALE |
| 9 | 1 | X1A.11 | NO DNA |  | 51 | 5 | X3B.04 | MALE |
| 10 | 1 | X1B.03 | NO DNA |  | 52 | 5 | X3A.03 | MALE |
| 11 | 1 | X1B.13 | NO DNA |  | 53 | 5 | X3A.11 | MALE |
| 12 | 1 | X1A.12 | NO DNA |  | 54 | 5 | X3B.03 | MALE |
| 13 | 1 | X1B.15 | NO DNA |  | 55 | 5 | X3B.15 | FEMALE |
| 14 | 1 | X1A.05 | NO DNA |  | 56 | 5 | X3A.28 | FEMALE |
| 15 | 1 | X1A.13 | NO DNA |  | 57 | 5 | X3A.25 | FEMALE |
| 16 | 1 | X2A.06 | NO DNA |  | 58 | 5 | X1B.05 | MALE |
| 17 | 1 | X1A.07 | NO DNA |  | 59 | 5 | X3B.01 | MALE |
| 18 | 1 | X1A.15 | NO DNA |  | 60 | 6 | X3B.02 | MALE |
| 19 | 1 | X1B.09 | NO DNA |  | 61 | 6 | X3B.19 | MALE |
| 20 | 1 | X1A.32 | NO DNA |  | 62 | 6 | X3A.31 | MALE |
| 21 | 1 | X2B.04 | NO DNA |  | 63 | 6 | X3A.09 | MALE |
| 22 | 1 | X1A.09 | NO DNA |  | 64 | 6 | X3B.06 | MALE |
| 23 | 1 | X1B.27 | FEMALE |  | 65 | 7 | X3B.20 | FEMALE |
| 24 | 1 | X1A.24 | MALE |  | 66 | 6 | X3A.23 | FEMALE |
| 25 | 1 | X2A.20 | MALE |  | 67 | 6 | X2B.09 | FEMALE |
| 26 | 1 | X2A.24 | MALE |  | 68 | 7 | X3B.30 | FEMALE |
| 27 | 1 | X1A.27 | MALE |  | 69 | 7 | X2B.19 | MALE |
| 28 | 1 | X1B.11 | NO DNA |  | 70 | 7 | X2A.25 | FEMALE |
| 29 | 1 | X2B.06 | FEMALE |  | 71 | 7 | X2A.08 | FEMALE |
| 30 | 2 | X2A.17 | FEMALE |  | 72 | 7 | X2A.03 | MALE |
| 31 | 2 | X2B.26 | FEMALE |  | 73 | 7 | X3A.30 | FEMALE |
| 32 | 2 | X2A.27 | MALE |  | 74 | 7 | X3B.31 | FEMALE |
| 33 | 2 | X2A.23 | FEMALE |  | 75 | 7 | X2A.19 | FEMALE |
| 34 | 2 | X2A.26 | MALE |  | 76 | 7 | X1B.10 | MALE |
| 35 | 2 | X2A.11 | MALE |  | 77 | 8 | X3B.11 | MALE |
| 36 | 2 | X2A.28 | FEMALE |  | 78 | 8 | X3A.12 | MALE |
| 37 | 2 | X2B.28 | FEMALE |  | 79 | 8 | X1A.31 | MALE |
| 38 | 2 | X2B.12 | MALE |  | 80 | 8 | X3B.29 | MALE |
| 39 | 2 | X2B.18 | MALE |  | 81 | 8 | X3B.26 | FEMALE |
| 40 | 2 | X2B.21 | MALE |  | 82 | 8 | X3A.07 | FEMALE |
| 41 | 3 | X2B.15 | FEMALE |  | 82 | 8 | X1B.12 | FEMALE |
| 42 | 1 | X1A.22 | NO DNA |  | 84 | 8 | X2B.22 | FEMALE |

Supplemental Table S3.

Transcripts with significantly different expression in male versus female embryos during the first 3h of development.

| FlybaseID | Gene symbol | Chr | Bias |  | FlybaseID | Gene symbol | Chr | Bias |
| --- | --- | --- | --- | --- | --- | --- | --- | --- |
| FBgn0264442 | ab | 2L | F |  | FBgn0030289 | GCS1 | X | F |
| FBgn0000117 | arm | X | F |  | FBgn0020300 | geko | 3L | F |
| FBgn0000137 | ase | X | F |  | FBgn0027287 | Gmap | X | F |
| FBgn0002069 | AspRS | 2R | F |  | FBgn0037376 | Hat1 | 3R | F |
| FBgn0030960 | Atg101 | X | F |  | FBgn0040318 | HGTX | 3L | F |
| FBgn0004862 | bap | 3R | F |  | FBgn0003997 | hid | 3L | F |
| FBgn0288686 | betaTub60D | 2R | F |  | FBgn0004828 | His3.3B | X | F |
| FBgn0050362 | boly | 2R | M |  | FBgn0027596 | Kank | 2R | F |
| FBgn0000229 | bsk | 2L | F |  | FBgn0261955 | kdn | X | F |
| FBgn0029957 | CG12155 | X | F |  | FBgn0013469 | klu | 3L | F |
| FBgn0030420 | pira | X | F |  | FBgn0267348 | LanB2 | 3L | F |
| FBgn0029771 | CG12730 | X | F |  | FBgn0262109 | lncRNA:CR42862 | 3L | M |
| FBgn0030868 | CG12986 | X | F |  | FBgn0267665 | lncRNA:CR46003 | 3L | M |
| FBgn0030807 | CG13000 | X | F |  | FBgn0261260 | mgl | X | F |
| FBgn0035186 | CG13912 | 3L | F |  | FBgn0036486 | Msh6 | 3L | M |
| FBgn0029931 | CG14427 | X | F |  | FBgn0005616 | msl-2 | 2L | M |
| FBgn0031632 | CG15628 | 2L | M |  | FBgn0030766 | mthl1 | X | F |
| FBgn0029990 | CG2233 | X | F |  | FBgn0002873 | mud | X | F |
| FBgn0023526 | CG2865 | X | F |  | FBgn0016684 | NaPi-T | 2R | F |
| FBgn0023529 | Grp170 | X | F |  | FBgn0030605 | ND-B18 | X | M |
| FBgn0028491 | CG2930 | X | M |  | FBgn0030500 | Ndc80 | X | F |
| FBgn0040373 | CG3038 | X | F |  | FBgn0260011 | NimC4 | 2L | F |
| FBgn0053494 | CG33494 | 3R | F |  | FBgn0015777 | nrv2 | 2L | F |
| FBgn0085322 | CG34293 | 3R | F |  | FBgn0039678 | Obp99a | 3R | F |
| FBgn0085430 | Dora | X | F |  | FBgn0015524 | otp | 2R | F |
| FBgn0034958 | CG3907 | 2R | M |  | FBgn0004394 | pdm2 | 2L | F |
| FBgn0037801 | CG3999 | 3R | F |  | FBgn0027621 | Pfrx | X | F |
| FBgn0025627 | CG4194 | X | F |  | FBgn0013725 | phyl | 2R | F |
| FBgn0259244 | CG42342 | 3R | F |  | FBgn0025739 | pon | X | F |
| FBgn0263776 | CG43693 | 3L | F |  | FBgn0283741 | prage | X | F |
| FBgn0264711 | CG43980 | 3L | F |  | FBgn0267507 | pre-rRNA:CR45847 | rDNA | M |
| FBgn0030796 | CG4829 | X | F |  | FBgn0083940 | RhoU | X | F |
| FBgn0031913 | CG5958 | 2L | F |  | FBgn0005631 | robo1 | 2R | F |
| FBgn0038407 | CG6126 | 3R | F |  | FBgn0003300 | run | X | F |
| FBgn0030990 | CG7556 | X | F |  | FBgn0003302 | rux | X | F |
| FBgn0031001 | CG7884 | X | F |  | FBgn0004170 | sc | X | F |
| FBgn0030863 | CG8188 | X | F |  | FBgn0025682 | scf | 3L | F |
| FBgn0030854 | CG8289 | X | F |  | FBgn0004880 | scrt | 3L | F |
| FBgn0034479 | CG8654 | 2R | F |  | FBgn0003345 | sd | X | F |
| FBgn0036381 | CG8745 | 3L | F |  | FBgn0261873 | sdt | X | F |
| FBgn0030514 | Mrgn1 | X | M |  | FBgn0003411 | sisA | X | F |
| FBgn0029504 | CHES-1-like | X | F |  | FBgn0026179 | siz | 3L | F |
| FBgn0263257 | Cngl | X | F |  | FBgn0039873 | Smvt | 3R | F |
| FBgn0030028 | Corp | X | F |  | FBgn0003463 | sog | X | F |
| FBgn0025641 | DAAM | X | M |  | FBgn0036411 | Sox21a | 3L | F |
| FBgn0020493 | Dad | 3R | F |  | FBgn0020378 | Sp1 | X | F |
| FBgn0263930 | dally | 3L | M |  | FBgn0260440 | spdo | 3R | F |
| FBgn0260635 | Diap1 | 3L | F |  | FBgn0037684 | Srr | 3R | F |
| FBgn0020307 | dve | 2R | F |  | FBgn0003459 | stwl | 3L | M |
| FBgn0260400 | elav | X | M |  | FBgn0263755 | Su(var)3-9 | 3R | M |
| FBgn0013953 | Esp | 3R | F |  | FBgn0040271 | Sulf1 | 3R | F |
| FBgn0038665 | euc | 3R | M |  | FBgn0264270 | Sxl | X | F |
| FBgn0000320 | eya | 2L | F |  | FBgn0017482 | T3dh | 2R | M |
| FBgn0000635 | Fas2 | X | F |  | FBgn0267001 | Ten-a | X | F |
| FBgn0004898 | fd96Cb | 3R | F |  | FBgn0019650 | toy | 4 | F |
| FBgn0030092 | fh | X | F |  | FBgn0046687 | Tre1 | X | F |
| FBgn0000658 | fj | 2R | F |  | FBgn0035521 | VhaM9.7-a | 3L | M |
| FBgn0037724 | Fst | 3R | M |  | FBgn0086680 | vvl | 3L | F |
| FBgn0016797 | fz2 | 3L | M |  | FBgn0010453 | Wnt4 | 2L | F |
| FBgn0038391 | GATAe | 3R | F |  | FBgn0001983 | wor | 2L | F |
| Bias indicates whether transcripts had higher expression in F (females) or higher expression in M (males) compared with the opposite sex at any point of the pseudo time. Chr, chromosome. | | | | | | | | |

Supplemental Table S4.

Source code for this manuscript.

#1.- Identification of unfertilized eggs.

library(RaceID)

samples <-read.csv("01raw_reads_n192samples.txt", sep="\t", header=TRUE, row.names = 1)

sc <- SCseq(samples)

sc<-filterdata(sc, minexpr = 3, minnumber = 5, LBatch = NULL, mintotal=250000)

sc <- compdist(sc,metric="spearman", FSelect = FALSE,knn = NULL,alpha = 3)

sc <- clustexp(sc, rseed = 12345, samp = 1000 , FUNcluster = "kmedoids", verbose = F)

sc <- findoutliers(sc, probthr = 0.001, outlg = 3, outminc = 5, verbose = TRUE)

pdf(file = "01results_maps.pdf", width = 7, height = 5)

sc <- comptsne(sc,perplexity = 16, rseed = 420)

sc <- compfr(sc,knn=10)

plotmap(sc,cex=3)

plotmap(sc,cex=3,fr=TRUE)

plotlabelsmap(sc, cex = 0.2)

plotexpmap(sc, g="scw", n="scw", logsc = TRUE, cex = 3)

plotexpmap(sc, g="sc", n="sc", logsc = TRUE, cex = 3)

plotexpmap(sc, g="esg", n="esg", logsc = TRUE, cex = 3)

types <- sub("(\\_\\d+)$","", colnames(sc@ndata))

subset <- types[grep("[A]",types)]

plotsymbolsmap(sc,types,subset=subset,cex=3,leg=F,

map=T, samples_col = rep("goldenrod1",400))

subset <- types[grep("[B]",types)]

plotsymbolsmap(sc,types,subset=subset,cex=3,leg=F,

map=T, samples_col = rep("orange",400))

dev.off()

#2.- Comparisons with previously published data and identification of older than 3h embryos.

library(RaceID)

library(RColorBrewer)

library(FateID)

samples <-read.csv("01raw_reads_n192samples.txt", sep="\t", header=TRUE, row.names = 1)

excluded <- c("X3A.19","X3A.14","X2A.29","X2A.16","X2A.12") #identified unfertilized eggs.

samples <- samples[,!(names(samples) %in% excluded)]

sc <- SCseq(samples)

sc<-filterdata(sc, minexpr = 3, minnumber = 5, LBatch = NULL, mintotal=250000)

sc <- compdist(sc,metric="spearman", FSelect = FALSE,knn = NULL,alpha = 3)

sc <- clustexp(sc, rseed = 12345, samp = 1000 , FUNcluster = "kmedoids", verbose = F)

sc <- findoutliers(sc, probthr = 0.001, outlg = 3, outminc = 5, verbose = TRUE)

pdf(file = "02results_maps_filtered.pdf", width = 7, height = 5)

sc <- comptsne(sc,perplexity = 15, rseed = 420)

sc <- compfr(sc,knn=10)

plotmap(sc,cex=3)

plotmap(sc,cex=3,fr=TRUE)

plotlabelsmap(sc, cex = 0.2)

plotexpmap(sc, g="scw", n="scw", logsc = TRUE, cex = 3)

plotexpmap(sc, g="sc", n="sc", logsc = TRUE, cex = 3)

plotexpmap(sc, g="esg", n="esg", logsc = TRUE, cex = 3)

types <- sub("(\\_\\d+)$","", colnames(sc@ndata))

subset <- types[grep("[A]",types)]

plotsymbolsmap(sc,types,subset=subset,cex=3,leg=F,

map=T, samples_col = rep("goldenrod1",400))

subset <- types[grep("[B]",types)]

plotsymbolsmap(sc,types,subset=subset,cex=3,leg=F,

map=T, samples_col = rep("orange",400))

dev.off()

clusters <- sc@cpart

write.csv(clusters, file = "02results_clusters_filtered.csv")

pdf(file = "02results_lineage_analysis_filtered.pdf", width = 7, height = 5)

ltr <- Ltree(sc)

ltr <- compentropy(ltr)

ltr <- projcells(ltr,cthr=1,nmode=T,knn=3)

ltr <- projback(ltr,pdishuf = 100, fast=FALSE, rseed=17000)

ltr <- lineagegraph(ltr)

ltr <- comppvalue(ltr,pthr=0.05, sensitive = T)

x <- compscore(ltr)

plotspantree(ltr,cex = 3,projections = T)

dev.off()

n <- cellsfromtree(ltr,c(2,8,9,5,7,13,3,6,1,4,12,10,11)) #select pseudo-temporal order vector from StemID.

x <- getfdata(ltr@sc)

fs <- filterset(x,n=n$f, minexpr = 0, minnumber = 0) #additional filtering and subsetting of gene expression

y	<- ltr@sc@cpart[n$f]

length(y)

fcol <- ltr@sc@fcol

write.table(y,"02results_pseudotime_names_filtered.csv",col.names=TRUE,sep=",",quote=FALSE)

pdf(file="02results_Kwasnieski_and_Sandler_filtered.pdf",width = 7, height = 5)

COL<-brewer.pal(n=12,name="Set3")

plotexpmap(sc, logsc = TRUE, cex = 3,

g=c("E(spl)m4-BFM","Elba3","l(1)sc","bnk","amos","dunk","nullo","sisA","sc",

"tsg","zen2","CG13000","CG13711","CG13712","CG14014","bard","CG15876",

"CG18269","CG6885","CG16813"),

n="List from Kwasnieski et al., 2019 (NC7-9)")

plotexpression(fs, y, n$f, col = fcol, alpha=.2, types=NULL, ylab = "Normalized Read Counts",

g=c("E(spl)m4-BFM","Elba3","l(1)sc","bnk","amos","dunk","nullo","sisA","sc",

"tsg","zen2","CG13000","CG13711","CG13712","CG14014","bard","CG15876",

"CG18269","CG6885","CG16813"),

cluster=FALSE, logsc = F,name = "List from Kwasnieski et al., 2019 (NC7-9)")

plotexpressionProfile(fs, y, n$f, col = COL, alpha=.2, lwd=5, ylim = c(0,0.1), ylab = "Normalized Expression",

g=c("E(spl)m4-BFM","Elba3","l(1)sc","bnk","amos","dunk","nullo","sisA","sc",

"tsg","zen2","CG13000","CG13711","CG13712","CG14014","bard","CG15876",

"CG18269","CG6885","CG16813"),

cluster=FALSE, name="List from Kwasnieski et al., 2019 (NC7-9)")

plotexpmap(sc, logsc = TRUE, cex = 3,

g=c("stumps", "ush", "Meltrin", "if", "ind", "aos", "ths",

"pnr", "spi", "pyr", "grh", "phm", "wntD", "tin",

"htl", "tup", "NetA"),

n="List from Sandler and Stathopoulos, 2016 - (14A to 14B)")

plotexpression(fs, y, n$f, col = fcol, alpha=.2, types=NULL, ylab = "Normalized Read Counts",

g=c("stumps", "ush", "Meltrin", "if", "ind", "aos", "ths",

"pnr", "spi", "pyr", "grh", "phm", "wntD", "tin",

"htl", "tup", "NetA"),

cluster=FALSE, logsc = F,name ="List from Sandler and Stathopoulos, 2016 - (14A to 14B)")

plotexpressionProfile(fs, y, n$f, col = COL, alpha=.2, lwd=5, ylim = c(0,0.07), ylab = "Normalized Expression",

g=c("stumps", "ush", "Meltrin", "if", "ind", "aos", "ths",

"pnr", "spi", "pyr", "grh", "phm", "wntD", "tin",

"htl", "tup", "NetA"),

cluster=FALSE, name="List from Sandler and Stathopoulos, 2016 - (14A to 14B)")

dev.off()

#3.- Generation of pseudo-time using only 3h embryos and differential expression analysis.

library(RaceID)

library(RColorBrewer)

library(FateID)

samples <-read.csv("01raw_reads_n192samples.txt", sep="\t", header=TRUE, row.names = 1)

excluded <- c("X3A.19","X3A.14","X2A.29","X2A.16","X2A.12", #identified unfertilized eggs.

"X1A.04","X1B.22","X3B.05","X3B.22","X3A.27", #cluster 3 (older than 3h)

"X1B.17","X3A.21","X1A.28","X1B.28","X3B.28", #cluster 6 (older than 3h)

"X1A.01", "X1B.24","X1B.29", "X1B.31","X3B.25","X3B.27", #cluster 1 (older than 3h)

"X1A.17","X1A.18","X1B.23","X2B.07","X3A.08", #cluster 4 (older than 3h)

"X2B.03","X3B.23", #cluster 12 (older than 3h)

"X2A.22","X3A.24","X1A.14","X2A.10","X2B.30", #cluster 10 (older than 3h)

"X2B.02", "X2B.25") #cluster 11 (older than 3h)

samples <- samples[,!(names(samples) %in% excluded)]

sc <- SCseq(samples)

sc<-filterdata(sc, minexpr = 3, minnumber = 5, LBatch = NULL, mintotal=250000)

sc <- compdist(sc,metric="spearman", FSelect = FALSE,knn = NULL,alpha = 3)

sc <- clustexp(sc, rseed = 12345, samp = 1000 , FUNcluster = "kmedoids", verbose = F)

sc <- findoutliers(sc, probthr = 0.001, outlg = 3, outminc = 5, verbose = TRUE)

pdf(file = "03results_maps_filtered_3h.pdf", width = 7, height = 5)

sc <- comptsne(sc,perplexity = 11.5, rseed = 1234) #13.659 #10.5 #11.45 #11.2

sc <- compfr(sc,knn=10)

plotmap(sc,cex=3)

plotmap(sc,cex=3,fr=TRUE)

plotlabelsmap(sc, cex = 0.2)

plotexpmap(sc, g="scw", n="scw", logsc = TRUE, cex = 3)

plotexpmap(sc, g="sc", n="sc", logsc = TRUE, cex = 3)

plotexpmap(sc, g="esg", n="esg", logsc = TRUE, cex = 3)

types <- sub("(\\_\\d+)$","", colnames(sc@ndata))

subset <- types[grep("[A]",types)]

plotsymbolsmap(sc,types,subset=subset,cex=3,leg=F, map=T, samples_col = rep("goldenrod1",400))

subset <- types[grep("[B]",types)]

plotsymbolsmap(sc,types,subset=subset,cex=3,leg=F, map=T, samples_col = rep("orange",400))

dev.off()

pdf(file = "03results_stemID_cell_projection_filtered_3h.pdf", width = 7, height = 5)

ltr <- Ltree(sc) #construct object used for StemID analysis

ltr <- compentropy(ltr) #calculation of the transcriptome entropy of each cell

ltr <- projcells(ltr,cthr=1,nmode=T,knn=3) #cell projection calculation,cthr=1,nmode=TRUE,knn=3

ltr<-projback(ltr,pdishuf = 100, fast=FALSE, rseed=17000)

ltr <- lineagegraph(ltr) #lineage tree is inferred

ltr <- comppvalue(ltr,pthr=0.05, sensitive = T) #p-values for the links calculation

x <- compscore(ltr)

plotspantree(ltr,cex = 3,projections = T)

dev.off()

n <- cellsfromtree(ltr,c(1,5,4,6,2,9,3,7)) #select pseudo-temporal order vector from StemID.

x <- getfdata(ltr@sc)

fs <- filterset(x,n=n$f, minexpr = 0, minnumber = 0) #additional filtering and subsetting of gene expression

y	<- ltr@sc@cpart[n$f]

length(y)

fcol <- ltr@sc@fcol

write.table(y,"03results_pseudotime_order_filtered_3h.csv",col.names=TRUE,sep=",",quote=FALSE)

yy <- as.data.frame(y) #y is an integer and I don't know how to get rownames from it so I convert it into a data frame

list_pseudotime <- row.names(yy)

norm_counts <- as.matrix(getfdata(sc))

norm_counts <- norm_counts[, list_pseudotime] #to get counts on pseudotemporal order

write.csv(norm_counts, file = "03results_fateid_counts_ps_norm_filtered_3h.csv")

A <- names(sc@cpart)[sc@cpart %in% c(1)]

B <- names(sc@cpart)[sc@cpart %in% c(5)]

x <- diffexpnb(sc@expdata,n=c(A,B),DESeq = TRUE, A=A, B=B , method = "per-condition")

write.table(x$res,"03diffexpnb_1_cl1and5_filtered_3h.xls",col.names=TRUE,sep="\t",quote=FALSE)

A <- names(sc@cpart)[sc@cpart %in% c(5)]

B <- names(sc@cpart)[sc@cpart %in% c(4)]

x <- diffexpnb(sc@expdata,n=c(A,B),DESeq = TRUE, A=A, B=B , method = "per-condition")

write.table(x$res,"03diffexpnb_2_cl5and4_filtered_3h.xls",col.names=TRUE,sep="\t",quote=FALSE)

A <- names(sc@cpart)[sc@cpart %in% c(4)]

B <- names(sc@cpart)[sc@cpart %in% c(6)]

x <- diffexpnb(sc@expdata,n=c(A,B),DESeq = TRUE, A=A, B=B , method = "per-condition")

write.table(x$res,"03diffexpnb_3_cl4and6_filtered_3h.xls",col.names=TRUE,sep="\t",quote=FALSE)

A <- names(sc@cpart)[sc@cpart %in% c(6)]

B <- names(sc@cpart)[sc@cpart %in% c(2)]

x <- diffexpnb(sc@expdata,n=c(A,B),DESeq = TRUE, A=A, B=B , method = "per-condition")

write.table(x$res,"03diffexpnb_4_cl6and2_filtered_3h.xls",col.names=TRUE,sep="\t",quote=FALSE)

A <- names(sc@cpart)[sc@cpart %in% c(1)]

B <- names(sc@cpart)[sc@cpart %in% c(2)]

x <- diffexpnb(sc@expdata,n=c(A,B),DESeq = TRUE, A=A, B=B , method = "per-condition")

write.table(x$res,"03diffexpnb_5_cl1and2_filtered_3h.xls",col.names=TRUE,sep="\t",quote=FALSE)

#4.- Differential expression analysis between males and females in 3h embryos. Input data can be found in Supplemental_Table_S6.xlsx

library(splineTimeR)

library(Biobase)

read.counts <-read.csv("normalized_reads.csv", sep=",", header=TRUE, row.names = 1)

psinfo_table <- read.csv("metadata.csv", sep=",", header=TRUE)

psnames <- psinfo_table[,1]

pssamples <- read.counts[,psnames]

row.names(psinfo_table) <- colnames(pssamples)

psinfo_table$SampleName <- factor(psinfo_table$SampleName)

psinfo_table$Treatment <- factor(psinfo_table$Treatment)

psinfo_table$Replicate <- factor(psinfo_table$Replicate)

all(rownames(psinfo_table) %in% colnames(pssamples))

all(rownames(psinfo_table) == colnames(pssamples))

phenoData <- new("AnnotatedDataFrame", data=psinfo_table)

minimalSet <- ExpressionSet(assayData=as.matrix(pssamples), phenoData = phenoData)

diffExprs <- splineDiffExprs(eSetObject = minimalSet, df = 7,

cutoff.adj.pVal = 0.01, reference = "MALE",

intercept = TRUE)

write.csv(diffExprs, file = "04results_7df_q0.01.csv")

Supplemental Table S5.

Primer sequences used in qPCR experiments and amplicon size in base pairs.

| FlybaseID or Chr location |  | Gene symbol |  | Forward sequence (Fwd \| 5’-3’)  Reverse sequence (Rev \| 5’-3’) |  | Amplicon size (bp) |
| --- | --- | --- | --- | --- | --- | --- |
| FBgn0040809 |  | CG13465 |  | Fwd \| GCAGATCATCAGTAACCAACCTG  Rev \| GAAACAAGAGCGGTTTCGGCA |  | 53 |
| FBgn0010433 |  | ato |  | Fwd \| AAACGATTCCAGCCTCAGCA  Rev \| TCATCGAACAAGGCGGAGTT |  | 175 |
| FBgn0010109 |  | dpn |  | Fwd \| ACGACGATTTCGACTGCTCC  Rev \| CCATAATCGGTTTGTTGGTCTTTCT |  | 117 |
| FBgn0001174 |  | halo |  | Fwd \| GCACTGCACTCTGGACACTC  Rev \| TGGAAAAAGTAGCTCGGCCA |  | 104 |
| FBgn0001090 |  | bnb |  | Fwd \| GTGCGTTGTGTGTTTTGTTGC  Rev \| GCAGTTGGCACTTCATTCTTTTG |  | 179 |
| FBgn0015949 |  | hrg |  | Fwd \| CGGCACCGACAATTTCTTTGTA  Rev \| AGGCATCAATGCCGAGGAAA |  | 101 |
| FBgn0004170 |  | sc |  | Fwd \| CCAACGACCATCAATTCGGC  Rev \| CGAGGAACCAGGCGATAGAG |  | 195 |
| FBgn0036058 |  | CG6707 |  | Fwd \| CAGAGTACCAACACCGGAGG  Rev \| GAGCGTTCCGAATGGGAGTA |  | 136 |
| FBgn0023517 |  | Pgam5 |  | Fwd \| GGCATCAAGTGGGACAAGGT  Rev \| CGTCGCGAAGGAAAGATGC |  | 190 |
| ChrX:4016153..4016261 |  | N/A |  | Fwd \| TTGGGCTGCTTCAGGTTTGA  Rev \| GAAGAGACACGCCAAGGCTA |  | 109 |
| ChrY:200524..200676 |  | N/A |  | Fwd \| TCATAAGGAGTGAAGCGGTCC  Rev \| AATTGTGTGCATCGGTGGGTC |  | 153 |
| All primers were designed and optimized in the lab. bp, base pairs; Chr, chromosome; N/A, not applicable | | | | | | |
